# Huntington’s disease onset is determined by length of uninterrupted CAG, not encoded polyglutamine, and is modified by DNA maintenance mechanisms

**DOI:** 10.1101/529768

**Authors:** Genetic Modifiers of Huntington’s Disease (GeM-HD) Consortium, James F. Gusella

## Abstract

The effects of variable, glutamine-encoding, CAA interruptions indicate that a property of the uninterrupted *HTT* CAG repeat sequence, distinct from huntingtin’s polyglutamine segment, dictates the rate at which HD develops. The timing of onset shows no significant association with *HTT cis*-eQTLs but is influenced, sometimes in a sex-specific manner, by polymorphic variation at multiple DNA maintenance genes, suggesting that the special onset-determining property of the uninterrupted CAG repeat is a propensity for length instability that leads to its somatic expansion. Additional naturally-occurring genetic modifier loci, defined by GWAS, may influence HD pathogenesis through other mechanisms. These findings have profound implications for the pathogenesis of HD and other repeat diseases and question a fundamental premise of the “polyglutamine disorders”.

## INTRODUCTION

Huntington’s disease (HD) is the most frequent of the polyglutamine diseases: dominantly inherited neurological disorders in which an expanded CAG trinucleotide repeat lengthens a segment of encoded glutamines in a particular protein (Lieberman et al., 2018). The distinct neuropathology and clinical manifestations and of each disorder are proposed to result from polyglutamine effects on the expression, activity and/or functional interactions of the encoded protein or of its polyglutamine-containing fragments. Nevertheless, in all, age-at-onset is inversely correlated with mutant CAG repeat size (Gusella and MacDonald, 2000). In HD, the expanded CAG tract lies in exon 1 of *HTT*, which encodes huntingtin, a large (>340 kDa), highly conserved, largely α-helical HEAT (huntingtin, elongation factor 3, protein phosphatase 2A and lipid kinase TOR) repeat protein (Guo et al., 2018). Neuronal loss in HD is most prominent in the striatum and other basal ganglia structures but also occurs throughout the cerebral cortex and in other regions. Post-mortem studies have demonstrated protein aggregates containing an N-terminal polyglutamine-containing fragment of mutant huntingtin in HD brain, suggesting a possible role for the formation of amyloid-like deposits in driving pathogenesis (Bates et al., 2015). Despite the decades of evidence that expressing long polyglutamine tracts in huntingtin disrupts various aspects of cellular function, HD pathogenesis is not well-understood and no disease-modifying treatments are available.

Current hopes for an effective intervention lie in suppressing expression of mutant huntingtin (Wild and Tabrizi, 2017), but subtle alterations in brain can be detected a decade or more before clinical onset (Paulsen et al., 2008). While the size of the expanded *HTT* CAG repeat explains ~60% of the individual variation in HD age-at-onset, the remaining variation shows heritability (Djousse et al., 2003). This prompted a human genetics strategy to identify disease-modifying factors that act prior to clinical diagnosis to either delay or hasten onset, with the expectation that, since they are validated in humans, these disease-modifying genes could reveal biochemical processes to target in drug development (GeM-HD Consortium, 2015). Our previous genome-wide association study (GWAS) of 4,082 individuals with HD, using the difference between age-at-onset predicted by CAG length and actual age-at-onset of motor symptoms, found three significant modifier signals at two loci, with pathway analysis suggesting DNA maintenance and mitochondrial regulation as potential modifying processes. We have now extended this GWAS strategy to over 9,000 HD individuals, increasing its power to detect both infrequent modifier alleles of strong effect and common modifiers with more modest impact. Our findings reveal effects of glutamine-encoding CAA interruptions within the *HTT* CAG repeat that question a fundamental premise of the polyglutamine diseases, as they indicate that the driver of the rate of HD pathogenesis leading to diagnosis is not the length of the polyglutamine tract in huntingtin, but rather the length of the uninterrupted CAG repeat segment in *HTT*. These findings cement the role of DNA maintenance mechanisms in modifying HD pathogenesis, most likely through an influence on somatic expansion of the *HTT* uninterrupted CAG repeat, and they disclose new loci that may influence HD pathogenesis through other mechanisms.

## RESULTS

### Genome-wide Association Analyses

Our previous GWAS (referred to here as GWA123) to identify genetic factors that influence age-at-onset involved independent groups of HD individuals genotyped in three stages (GWA1, GWA2 and GWA3) and analyzed using a robust statistical phenotype model relating the CAG repeat length to log-transformed age-at-onset of diagnostic motor signs of HD individuals with 40–55 CAG repeats (GeM-HD Consortium, 2015). This regression model yielded a residual age-at-onset value for each subject that was transformed back into natural scale as a phenotype for continuous, quantitative association analysis. We used the same approach here (Figure S1A) in additional European-ancestry individuals with HD from Registry (N=4,986) (Orth et al., 2010) and from the Enroll-HD platform of the CHDI Foundation (N=1,312) (Landwehrmeyer et al., 2017), genotyped in separate waves (GWA4 and GWA5, respectively). The distribution of residuals for the combined GWA12345 dataset was similar to a theoretical normal distribution (Figure S1B). In view of improvements in reference databases and imputation, we re-imputed GWA123 to match the imputation of GWA45 using the Haplotype Reference Consortium (HRC) (McCarthy et al., 2016). We then analyzed the 9,064 unique individuals (4,417 males; 4,647 females) in the combined GWA12345 dataset using two parallel approaches: continuous analysis of association to residual age-at-onset and dichotomous analysis of extremes of residual age-at-onset (Figure 1 and Table 1). The former strategy is more effective for detecting rarer modifier alleles whereas the latter is less influenced by slight imprecision in establishing age-at-onset since it compares overall allele frequencies between groups who have onset substantially later or earlier than expected. For comparison, results from the combined GWA45 dataset alone and from all GWA datasets subjected to meta-analysis are shown in Figure S1C&D along with Q-Q plots for the overall GWA12345 continuous and dichotomous analyses in Figure S1E&F.

**Table 1:**
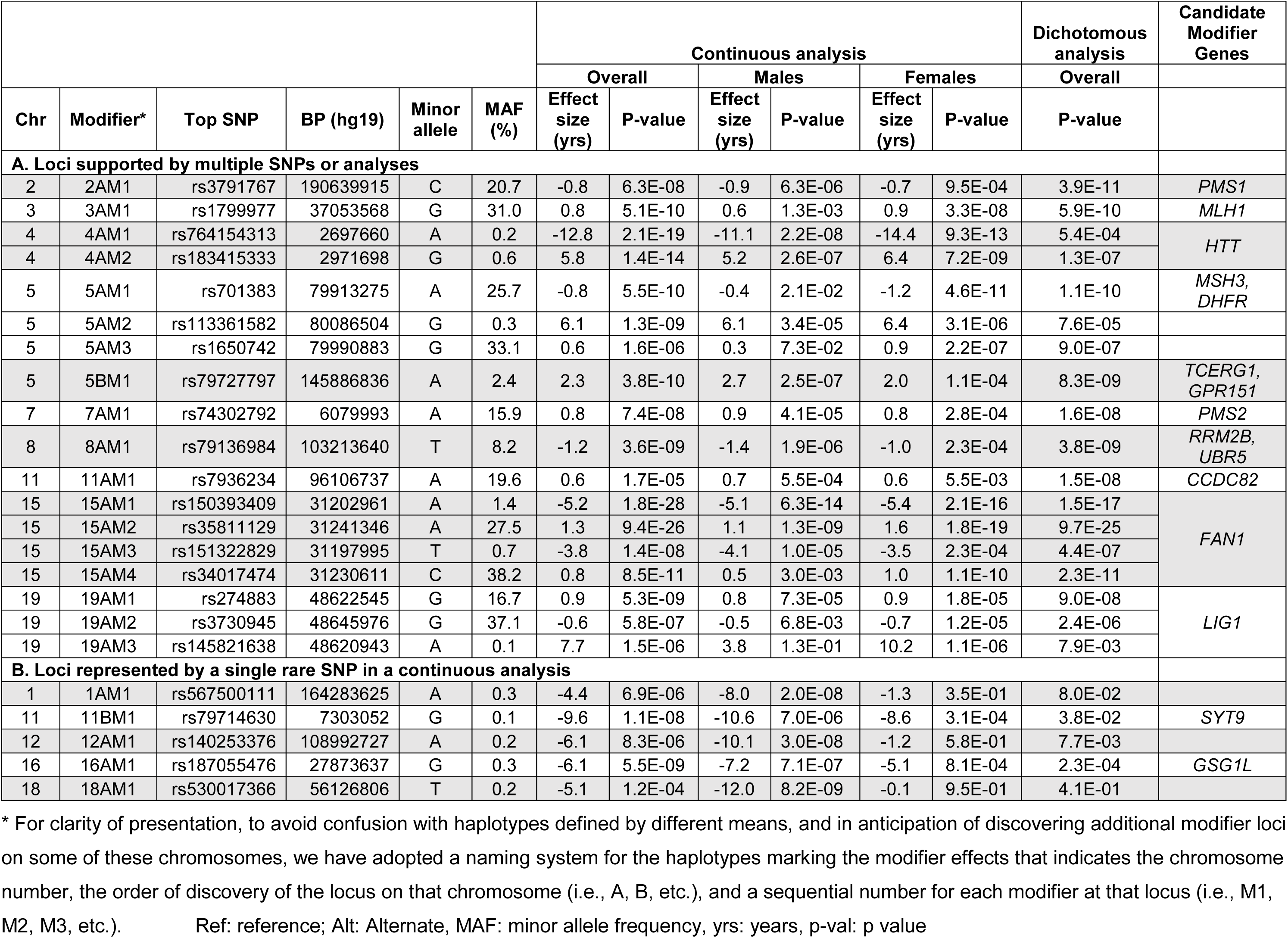
Genome-wide Significant Loci with Additional Modifier Haplotypes Identified by Conditional Analysis

**Figure 1.**
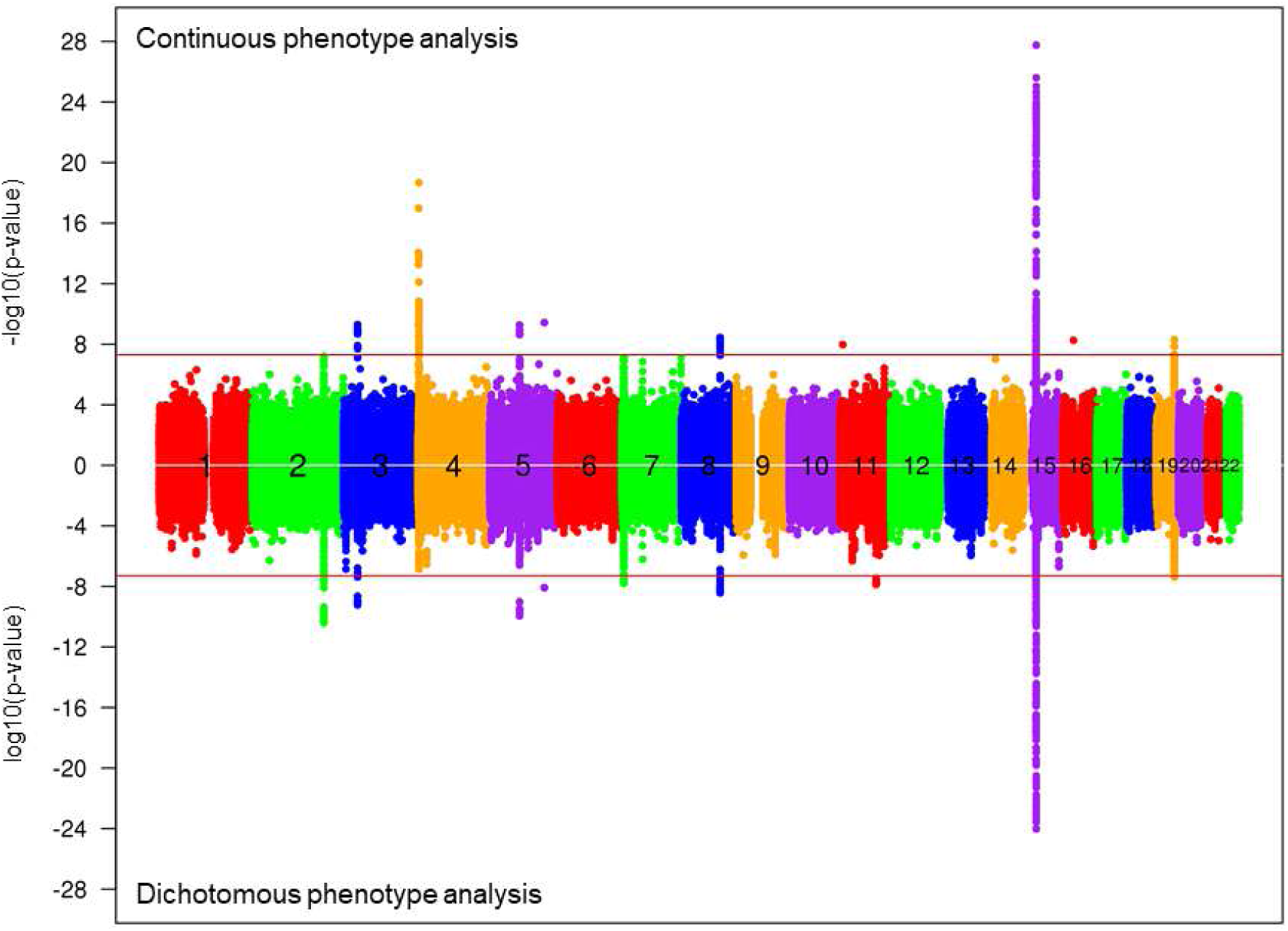
Manhattan plots showing HD onset modification signals. Results of GWA analysis using the continuous residual age-at-onset phenotype or a dichotomized phenotype are summarized, with each circle representing a test SNP and significances represented as -log10(p-value) (top) and log10(p-value) (bottom), respectively.

In GWA123, we had observed modifier loci on chromosome (chr) 8 and chr 15, with the latter exhibiting two independent opposing effects associated with different tag SNPs (GeM-HD Consortium, 2015). In follow-up studies, a chr 3 locus also achieved genome-wide significance (Lee et al., 2017). These loci emerged again in GWA12345, along with several new loci genome-wide significant in either continuous or dichotomous analysis or both. Like the chr3 (*MLH1*) and chr15 (*FAN1*) loci, the new loci on chr 2 (*PMS1*), 5 (*MSH3*), 7 (*PMS2*) and 19 (*LIG1*) all contained genes associated with aspects of DNA repair but additional modifier sites on chr 5 (*TCERG1, GPR151*) and chr 11 (*CCDC82*) may not be directly connected to these processes. A robust association signal was also obtained on chr 4, in the vicinity of *HTT*. Three of these new loci, on chr 4, 5 and 19, showed evidence (p <1E-5) upon conditional analysis of more than one independent modifier effect tagged by different SNPs while the chr 15 locus revealed 2 additional modifier effects. Two additional genome-wide significant loci, on chr 11 (*SYT9*) and chr 16 (*GSG1L*), displayed signal only in the continuous analysis from a single very low-frequency SNP allele, leaving open the possibility of a statistical artifact due to extreme phenotypic outliers. A larger sample size and/or functional analysis will be needed to firmly establish these loci as *bona fide* modifiers.

Although there was no significant difference in age-at-onset of HD between the sexes (Figure S2A), we also performed male- and female-specific association analyses, which revealed differences in relative effect size and significance of association for some modifiers (Figure S2 B-E and Table 1). This was most evident for the chr 5 (*MSH3*) locus, where the common modifiers had a far greater effect in females. The sex specific analysis also revealed 3 new loci (chr 1, 12 and 18) that may contain male-specific modifiers or, since they are tagged by very rare alleles, may be due to statistical outlier effects.

A list of all SNPs yielding suggestive signals (p <1E-5) in any of the single SNP association analyses is given in Table S1. We also performed gene-wide association analysis (Table S2) using a window spanning 35kb upstream and 10kb downstream of the transcript (to capture regulatory elements) (Network and Pathway Analysis Subgroup of Psychiatric Genomics, 2015). This yielded respectively 45 and 12 genome-wide significant genes in GWA12345 and GWA45 alone, of which 34 and 11 were significant when restricting analysis to the transcript boundaries. All significant genes are from regions in Table 1A. Analysis of pathway enrichment in both GWA12345 and GWA45, compared to GWA123, again pointed to mismatch repair, while mitochondrial regulation was no longer supported (Table S3). In 77 gene sets related to various aspects of DNA repair (Pearl et al., 2015), the strongest enrichments were in gene sets related to mismatch repair (Table S4). The most significant genes are shown in Table S5. In a broader test of 14,210 pathways with 10 or more gene members, 77 pathways were significant after Bonferroni correction in GWA12345 (Table S6). The top 13 pathways were all related to DNA maintenance processes, but some top GWAS genes, such as *FAN1, RRM2B* and *UBR5*, appeared only in the largest, most general pathway (GO 6281: DNA repair). Taken together, these analyses provide robust evidence for genes involved in DNA mismatch repair pathways in the genetic modification of HD age-at-onset but suggest that other DNA maintenance processes and potentially non-DNA-related processes also play a role.

We next assessed potential association of gene expression and residual age-at-onset by performing a transcriptome-wide association study (TWAS), imputing the Common Mind Consortium prefrontal cortex expression to the GWA12345 summary association statistics (Gusev et al., 2016). Four genes at 3 loci (*FAN1* on chr 15: 1.86E-22, *MSH3* on chr 5: 1.90E-8, and *PMS1*: 6.11E-7 and *ASNSD1*: 5.26E-6, both on chr 2) were significant after Bonferroni correction, with increased *FAN1*, *PMS1*, and *ASNSD1* expression and decreased *MSH3* expression associated with later onset. Conditioning on expression removed most of the association signal for *MSH3* and *PMS1/ASNSD1*, while also removing signal corresponding to the common onset-delaying, but not the rarer onset-hastening, GWAS SNPs at *FAN1*.

Comparison of the HESS regional heritability estimates (Shi et al., 2016) before and after conditioning on expression predict that 40%, 57% and 87% of the contribution of *FAN1*, *PMS1* and *MSH3*, respectively, to the genetic age-at-onset liability can be explained by *cis* expression effects.

### The ostensible chromosome 4 ‘modifier’ locus - *HTT*

It was surprising that GWA12345 showed a clear signal on chr 4 in the vicinity of *HTT* as we have shown previously that in individuals with HD, the most frequent 7 *HTT* SNP haplotypes with CAG expansions on disease chromosomes are not associated with differences in age-at-onset (Lee et al., 2012a). In addition, there is no modifier effect either of *HTT* haplotype or CAG repeat size on the normal chromosome in HD heterozygotes (Lee et al., 2012b). Upon conditional analysis (Figure 2A) the chr 4 GWAS signal resolved into two independent effects tagged by rare alleles near *HTT*, an onset-hastening effect of ~12.7 years per allele (haplotype 4AM1) and an onset-delaying effect of ~5.7 years per allele (haplotype 4AM2).

**Figure 2.**
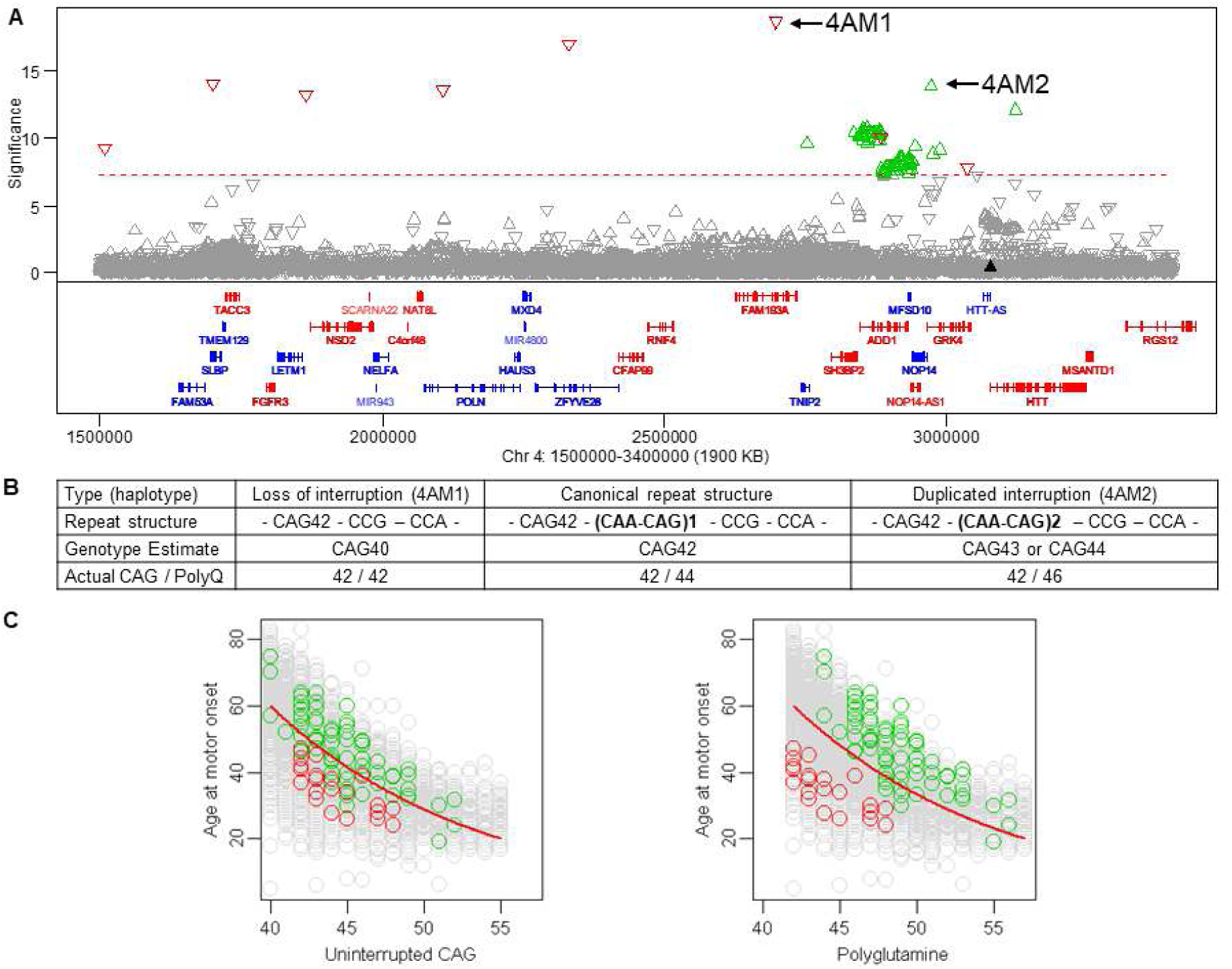
Ostensible onset modification signals on chromosome 4. **A**. Signals from the chr 4 region of significant association (continuous phenotype) are plotted against genomic coordinate (GRCh37/hg19) with a dotted red line indicating genome-wide significance and red and green triangles representing independent onset-hastening (4AM1) and onset-delaying (4AM2) haplotypes, respectively. In this and subsequent plots, downward and upward triangles represent SNPs whose minor alleles are associated with hastened and delayed onset, respectively. Genes are displayed below in red (plus strand) and blue (minus strand), with all RefSeq exons combined to plot exon-intron structures. **B**. *HTT* repeat structures, genotyping estimates, lengths of uninterrupted CAG, and lengths of polyglutamine for 4AM1 (left), canonical (center), and 4AM2 haplotypes are illustrated using uninterrupted CAG42 as an example. **C**. MiSeq analysis of 4AM1 (red circles) and 4AM2 (green circles) individuals identified those with non-canonical repeats, permitting comparison of their age-at-onset with uninterrupted CAG repeat size (left panel) or polyglutamine length (right panel). Red trend lines represent our standard onset-CAG phenotyping model. Grey circles represent HD individuals who do not carry 4AM1 or 4AM2 modifier haplotype. Of 29 individuals with the 4AM1 tag SNP, 28 had DNA available for sequencing: 21 with no CAA on the disease chr (mean residual (mr) −15.96), 2 with no CAA on the normal chr (mr 4.77) and 5 with canonical sequence on both (mr −3.07). Of 102 individuals with the 4AM2 tag SNP, 98 had DNA available: 68 with a second CAA on the disease chr (mr 7.21), 23 with a second CAA on the normal chr (mr 1.96) and 7 canonical on both (mr 1.65).

Recently, it was proposed that an *HTT* promotor SNP, rs13102260, alters HD onset through an effect on *HTT* expression on both normal and disease chromosomes (Becanovic et al., 2015), so we first examined the possibility that genetic effects on *HTT* mRNA production were responsible for the apparent modifier signals. In normal brain data from the GTEx Consortium, *HTT* mRNA showed wide variation in expression (Figure S3A) but significant *cis*-eQTL SNPs in the GTEx brain regions showed no correspondence with the age-at-onset association signals in our GWAS (Figure S3C-F). Notably, rs13102260 was not significant (black triangle - Figure 2A & S3C-F). In addition, individuals with two CAG alleles in the expanded range (and therefore no normal *HTT* allele) who express twice the level of mutant huntingtin have age-at-onset residuals (based upon the longer of the two expanded repeats) that do not differ from heterozygotes with only one copy of the disease gene (Figure S3B). Taken together, these findings suggest that variation in *HTT* mRNA expression within the normal physiological range does not significantly influence HD age-at-onset.

### Uninterrupted CAG repeat rather than polyglutamine determines HD onset

The vast majority (>95%) of European ancestry HD chromosomes carry a canonical sequence of the CAG repeat tract that includes a penultimate CAA interruption followed by a CAG at the 3’ end of the repeat (Figure 2B, Table S7). Since both CAA and CAG encode glutamine, the length of the polyglutamine segment derived from this canonical structure is consistently greater than the length of the uninterrupted CAG tract by 2 residues, so it has not been possible to distinguish whether the correlation with age-at-onset is due to polyglutamine size or to CAG repeat size. For GWA12345, the uninterrupted CAG repeat length was estimated by a PCR fragment-sizing assay in comparison to uninterrupted CAG repeat length in sequenced assay DNA standards with this canonical sequence. We reasoned that if haplotypes 4AM1 and 4AM2 carried rare non-canonical sequence variations, an inaccurate uninterrupted CAG repeat estimation from the standard assay may artificially affect residual age-at-onset, thereby resulting in the apparent modifier effects. Indeed, sequencing of the *HTT* CAG repeat region revealed that the onset-hastening 4AM1 haplotype was associated with the loss of the canonical penultimate CAA interruption (Figure 2B) such that the PCR genotyping assay underestimated the length of the uninterrupted CAG repeat by 2 CAGs. Conversely, the onset-delaying 4AM2 haplotype was associated with a second CAA interruption, two codons upstream of the first. The resulting overestimation of the uninterrupted CAG repeat length was of 1 or 2 residues rather than always 2 residues, due to mis-priming of the PCR primer on this sequence variation in the genotyping assay. When we used the true uninterrupted CAG length, both association signals were very substantially less significant (4AM1: 12.83E-8 from 2.12E-19; 4AM2: 1.71E-5 from 1.42E-14), indicating that most of the apparent modifier signal was due to the difference in age-at-onset normally expected for a CAG repeat length mis-estimated by 1-2 CAGs.

Importantly, 4AM1 and 4AM2 haplotypes allow the role of the uninterrupted CAG repeat size to be distinguished from that of polyglutamine in determining age-at-onset. The 4AM1 repeat sequence variation encodes the same number of glutamines as consecutive CAGs while that of 4AM2 specifies 4 more glutamines than consecutive CAGs (Figure 2B). As demonstrated in Figure 2C, age-at-onset tracks best with the length of the uninterrupted CAG repeat in *HTT* and is not equivalently influenced by the length of the polyglutamine segment in huntingtin.

Furthermore, the much reduced but still significant association signals remaining unaccounted for after correcting the uninterrupted CAG repeat length for 4AM1 and 4AM2 imply functional differences from canonical sequence haplotypes. These effects are not due to 4AM1 or 4AM2 haplotypes on normal chromosomes since removing only individuals with the tag SNPs on the disease chromosome eliminated the association signal. Neither is polyglutamine toxicity involved, since for any matching uninterrupted CAG size, 4AM2 has 4 more glutamines than 4AM1 and yet is associated with later rather than earlier onset. From the previous eQTL analyses, *HTT* mRNA levels are not likely to be responsible, although an influence of the interruptions on translation is formally possible. Similarly, an additional haplotype-specific influence on HD acting through another gene distal to *HTT* is conceivable, but Occam’s razor suggests a more likely explanation. The critical importance of a DNA sequence property rather than polyglutamine length in determining HD onset suggests that the DNA sequence context in which the CAG repeat is embedded also has an influence. On 4AM1 HD chromosomes, the ^CAG repeat is followed by CCG_12_CCT_2_CAGCTTCCT_1_ and on 4AM2 chromosomes, it is followed by CAA_1_CAG_1_CAA_1_CAG_1_CCG_1_CCA_1_CCG_7_CCT_3_CAGCTTCCT_1_. Sequencing of samples not^ tagged by 4AM1 and 4AM2 SNPs confirmed correct PCR-assay genotyping of the uninterrupted CAG sequence on canonical chromosomes and showed that the vast majority carry CAA_1_CAG_1_CCG_1_CCA_1_CCG_7_CCT_2_CAGCTTCCT_1_ after the CAG repeat (Table S7). An influence of the downstream sequences on the onset-determining property of the uninterrupted CAG repeat would both explain the remaining signal and argue that other sequence variations on even less frequent HD chromosomes might also influence age-at-onset but not be detected with the current GWAS sample size. Indeed, in addition to those carrying 4AM1 or 4AM2 tag SNPs, the sequences of a selection of both HD and normal chromosomes revealed a wide variety of rarer non-canonical sequence variations that could potentially influence the critical property of the adjacent uninterrupted CAG repeat (Table S7).

### A chromosome 5 modifier locus – *MSH3 / DHFR*

The most prominent modifier locus on chr 5 centers on *MSH3*, which encodes a DNA mismatch repair protein, and *DHFR*, which produces dihydrofolate reductase, a critical enzyme in determining nucleotide pools. These genes are notable for sharing a bidirectional promotor (Shinya and Shimada, 1994). The top SNP tagged a relatively frequent onset-hastening modifier effect (haplotype 5AM1). Conditional analyses revealed 2 additional onset-delaying haplotypes (5AM2 and 5AM3), tagged by relatively infrequent and more common alleles, respectively (Figure 3, Table 1). This locus has previously been implicated as harboring a modifier that influences a multi-factor measure of progression of HD deterioration (Moss et al., 2017) but we could not directly assess that tag SNP, reported as a missense change within *MSH3*, because the variant maps within a set of tandem repeats and is not imputed using HRC data. However, using the 1000 Genomes as a reference set (1000 Genomes Project Consortium, 2015), we re-examined the conditional analysis and found that the haplotype reported as a progression modifier in TRACK-HD is equivalent to 5AM3.

**Figure 3.**
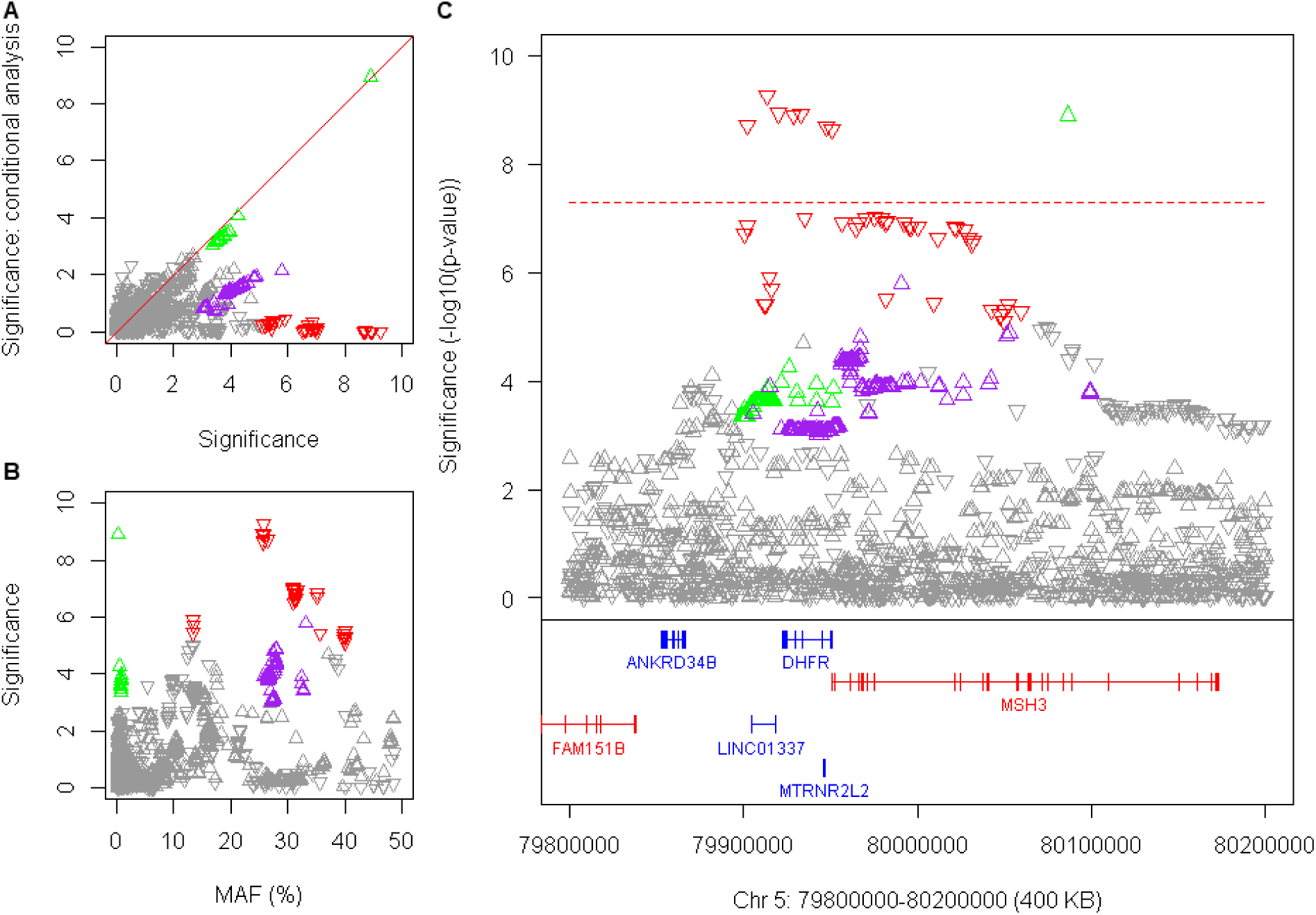
HD onset modification signals at *DHFR / MSH3* on chromosome 5. **A**. Significance of SNP association in continuous analysis conditioned on the top SNP in the region (rs701383, onset-hastening haplotype 5AM1, red downward triangles) is compared to GWA12345 significance to detect independent modifier haplotypes with greater than suggestive significance (p-value < 1E-5): 5AM2 (green) and 5AM3 (purple), onset-delaying (upward triangles). **B**. The significance of SNP association in GWA12345 is compared to minor allele frequency. **C**. Significance for SNPs from the three independent modifier haplotypes is shown relative to genes in the region.

In GTEx Consortium data, the top SNP alleles from 5AM1 corresponded strongly to *cis-*eQTLs associated with increased expression of *MSH3* (but not *DHFR*) in blood (Figure S4A&B). In mice, several mismatch repair genes, including *Msh3*, have been reported to influence the somatic instability of CAG repeats (Dragileva et al., 2009; Schmidt and Pearson, 2016; Tome et al., 2013) and in humans, naturally occurring *MSH3* polymorphisms were associated with instability of the non-coding myotonic dystrophy type 1 CTG repeat in blood (Morales et al., 2016). Based on data from *Msh3* knock-out mouse models, where somatic CAG expansion was suppressed, the onset-hastening 5AM1 associated with increased *MSH3* expression would be predicted to be associated with higher levels of somatic CAG expansion, a hypothesis that we were able to test using ABI GeneMapper fragment sizing expanded allele profiles from our CAG genotyping assay. The bulk of the PCR product for any individual corresponds to the presumptive inherited expanded CAG size and constitutes a floor with respect to which expansion can be examined. Those individuals with mosaicism for the highest somatic CAG expansions compared with this inherited size are evident from the increased fraction of larger PCR products detected. Among the 7,013 individuals with suitable traces, there was an inherent increase in expansion with CAG repeat size (Figure S4C) so we identified the 25% (N=1,753) who showed the highest proportion of somatic expansions at each repeat length from 40-55 and examined their genotype at rs701383. This 5AM1 tag SNP deviated significantly from Hardy-Weinberg expectation (chi-square 15.80, 2 d.f., p < 0.0004) due to an excess of the minor A allele (observed 915 GG, 684 AG, 154 GG vs. expected 968 GG, 669 AG, 116 AA) indicating that increased *MSH3* expression can be associated with increased somatic expansion. In the brain, the top 5AM1 SNPs from the GWAS correspond less well than in the blood with *MSH3 cis*-eQTL signals, suggesting that additional factors regulating the locus beyond the steady state may be important for determining CAG instability (Figure S4D). These additional influences may explain the sex difference in the effect of this modifier, since the *cis*-eQTLs do not appear to be sex specific (Figure S4 legend).

In contrast, the rare haplotype 5AM2 SNP alleles associated with delayed onset did not overlap with *cis*-eQTLs for either *MSH3* or *DHFR* (Figure S4D&E) and did not differ in effect size between the sexes, suggesting that this haplotype may harbor a variant that alters the mRNA and/or protein. SNP alleles tagging 5AM3, which has been connected both to delayed HD onset and to delayed progression, corresponded more robustly with *cis*-eQTLs for decreased *DHFR* expression in cortex, caudate, putamen and blood than with *MSH3 cis*-eQTLs (Figure S4D&E), suggesting potential roles for both *MSH3* and *DHFR* in HD modification. Although the 5AM3 tag SNPs and blood *cis*-eQTL SNPs correspond, if this onset-delaying modifier acted through DNA maintenance mechanisms, it would be expected to reduce the degree of CAG somatic expansion, a hypothesis that is not readily tested using our assay for extreme somatic expansions.

### Other Mismatch Repair Genes – *MLH1, PMS1, PMS2*

Three other mismatch repair genes were implicated by genome-wide significant signals in GWA12345. The chr 3 locus containing *MLH1* displayed a single onset-delaying modifier signal and we obtained no strong evidence in our TWAS or GTEx Consortium data analyses that this effect acts by altering gene expression. The peak SNP, rs1799977, specifies a missense change, I219V, that is considered benign (SIFT: tolerated; Polyphen: benign) but this does not exclude a subtle effect on the activity or interactions of MLH1 in the context of CAG repeat expansion. The genes *PMS1* and *PMS2* (postmeiotic segregation increased 1 and 2, respectively) both encode proteins that form heterodimers with MLH1 in the mismatch repair process (Pearl et al., 2015). *PMS1* and *PMS2* map to GWA12345 modifier peaks on chr 2 and chr 7, respectively. Upon conditional analysis, the chr 2 locus showed a single onset-hastening modifier effect and the chr 7 locus a single onset-delaying modifier effect. The TWAS evidence suggested that the chr 2 modifier may act through decreased expression of *PMS1*, at least in cortex, but there is no comparable expression evidence for *PMS2*. *PMS1* did not show strong *cis*-eQTLs in blood and the *PMS2* associated modifier delayed onset, so neither was testable for an effect of expression levels using our assay for extreme somatic expansions in that tissue. While we have not identified the functional variation at either locus and it remains formally possible that the modifier effect acts through another gene in each region, the most parsimonious explanation is that modification acts through these DNA repair genes.

### The chromosome 15 modifier locus – *FAN1*

The expanded GWA12345 analysis replicated at the chr 15 locus the opposing effects of an infrequent onset-hastening modifier (15AM1) and a frequent onset-delaying modifier (15AM2), revealed two novel modifier haplotypes, 15AM3 and 15AM4 (Figure 4, Table 1) and pointed to genetic variation affecting *FAN1* as the source of HD modification. *FAN1* encodes a nuclease with functions that include involvement in interstrand DNA crosslink (ICL) repair (Smogorzewska et al., 2010). It is also recruited to stalled replication forks, physically interacts with the mismatch repair protein MLH1, another of the proposed HD age-at-onset modifiers, and is needed for homologous recombination but not for double-strand break resection (Cannavo et al., 2007; Lachaud et al., 2016; MacKay et al., 2010). The peak SNP, tagging the onset-hastening 15AM1, is missense variant rs150393409 (R507H; SIFT: deleterious, Polyphen: benign) that has recently been associated in human population studies with karyomegalic interstitial nephritis, a recessively inherited disease caused by loss of FAN1 function (Bastarache et al., 2018). A second, less frequent missense variant, rs151322829 (R377W) tags the onset-hastening 15AM3 and is also predicted to be deleterious to FAN1 function by SIFT (SIFT: deleterious, Polyphen2: benign). Conversely, tag SNPs for the onset-delaying 15AM2 and 15AM4 correspond with *cis*-eQTL SNPs for increased expression in cortex in GTEx Consortium data (Figure S5A) and 15AM4 also matches such *cis*-eQTLs SNPs in blood (Figure S5B). Since the two onset-hastening haplotypes are both of low frequency neither they, nor the onset-delaying haplotypes, could be tested for hypothesized effects on CAG expansion in blood using our assay for somatic expansion. No other gene in the region shows a missense variant or correspondence between any cortex, caudate or putamen *cis*-eQTLs and modifier association signals, pointing like our TWAS findings to *FAN1* as the source of the modifier effects, with reduced function leading to hastened onset and increased expression leading to delayed onset. Notably, lowering *FAN1* expression in mammalian cells and patient-derived iPSCs induced *HTT* CAG expansions (Goold et al., 2018) and preliminary findings have indicated a similar effect of FAN1 deficiency on the CAG repeat in an HD knock-in mouse model (Loupe and MacDonald, unpublished results). In a mouse model of Fragile X syndrome, inactivation of *Fan1* induced somatic expansion of a CGG repeat (Zhao and Usdin, 2018) indicating that the impact of *FAN1* variation may extend to other non-CAG, non-coding repeat diseases.

**Figure 4.**
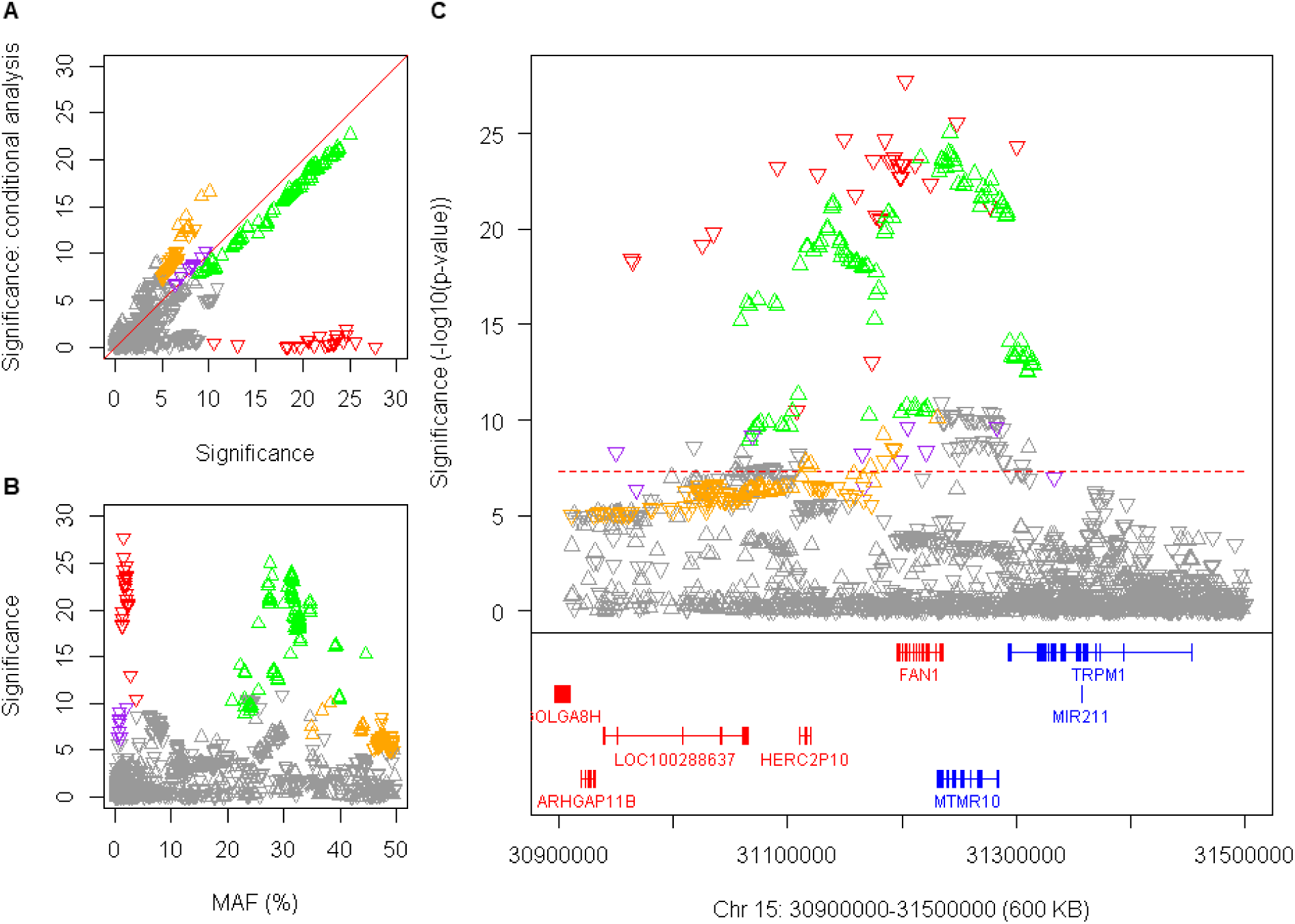
Four onset modification signals on chromosome 15. **A.** Significance of SNP association in continuous analysis conditioned on the top SNP in the region (rs150393409, onset-hastening haplotype 15AM1, red downward triangles) is compared to GWA12345 significance to detect independent modifier haplotypes with greater than suggestive significance (p-value < 1E-5): 15AM2 (green, onset-delaying), 15AM3 (purple, onset-hastening), and 15AM4 (gold, onset-delaying). Because SNPs tagging 15AM4 have alleles of close to equal frequency, the direction of the arrow varies depending on whether the minor or major allele is on the 15AM4 haplotype. **B**. The significance of SNP association in GWA12345 is compared to minor allele frequency. **C**. Significance for SNPs from the four independent modifier haplotypes is shown relative to genes in the region.

### The chromosome 19 modifier locus – *LIG1*

Another new modifier signal in GWA12345 mapped on chr 19 to *LIG1*, which encodes an ATP-dependent DNA ligase that seals DNA nicks during replication, recombination and a variety of DNA damage responses, including base-excision repair (Howes and Tomkinson, 2012). Conditional analysis revealed 3 different modifier haplotypes (Figure 5, Table 1). Common modifier haplotypes delayed (19AM1) or hastened (19AM2) onset by less than 1 year per allele, but a third modifier haplotype (19AM3) tagged by a rare variant had a strong effect on onset, delaying it by ~7.7 years. The top 19AM3 SNP, rs145821638, causes a missense change in LIG1 (K845N) that is predicted to be potentially damaging to its function (SIFT: deleterious; Polyphen: possibly damaging), suggesting that reduced activity and/or altered interactions of LIG1 protein may suppress CAG repeat expansion. Conversely, the onset-hastening haplotype 19AM2 is associated with increased expression of LIG1 in cortex and BA9 (Figure S6A), consistent with the increased CAG repeat instability observed due to exogenous overexpression of LIG1 in human cells (Lopez Castel et al., 2009). The mechanism by which the common but weaker onset-delaying 19AM1 might have its effect is not yet clear. None of the haplotype association signals showed correspondence with *cis*-eQTLs in blood (Figure S6B).

**Figure 5.**
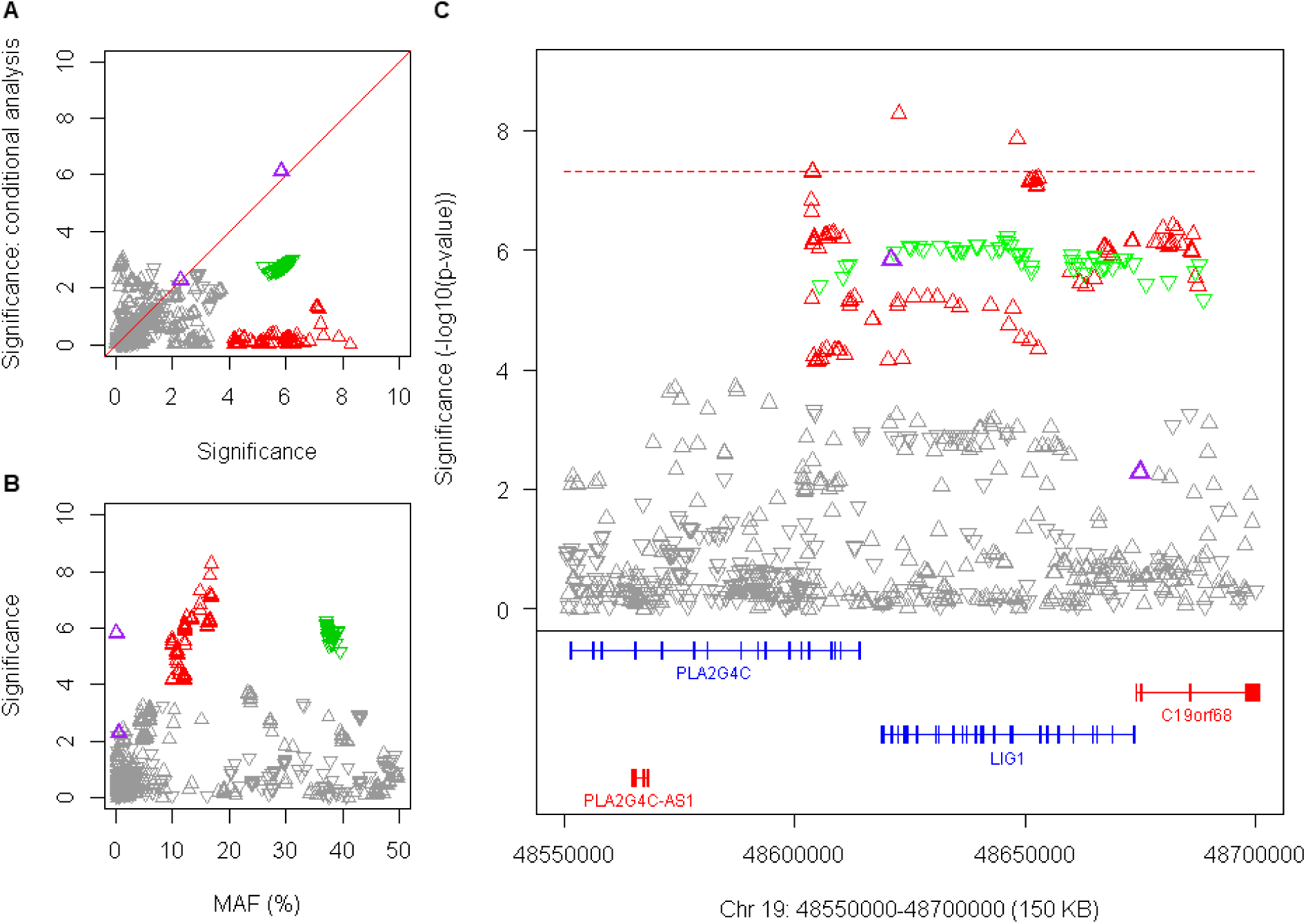
Three onset modification signals on chromosome 19. **A**. Significance of SNP association in continuous analysis conditioned on the top SNP in the region (rs274883, onset-delaying, 19AM1, red upward triangles) is compared to GWA12345 significance to detect independent modifier haplotypes with greater than suggestive significance (p-value < 1E-5): 19AM2 (green, onset-hastening) and 19AM3 (purple, onset-delaying). **B**. The significance of SNP association in GWA12345 is compared to minor allele frequency. **C**. Significance for SNPs from the three independent modifier haplotypes is shown relative to genes in the region.

### Individual GWAS loci that may reveal other mechanisms of modification

Any of the other modifier loci could theoretically act by an indirect impact on the same DNA maintenance processes that involve the above genes, or they could represent independent mechanisms of modification. It is not yet clear whether *RRM2B*, encoding the small subunit of a p53-inducible ribonucleotide reductase, or *UBR5*, specifying an E3 ubiquitin-protein ligase, mediates the modifier effect at the chr 8 locus and whether modification acts via influencing DNA maintenance (e.g., through regulating nucleotide pools, or levels of repair enzymes) or through some other mechanism. The chr 11 locus containing *CCDC82*, which encodes a coiled-coil domain protein, also includes *JRKL* (jerky homolog-like) specifying a likely nuclear regulator and a portion of *MAML2* (mastermind-like 2) which produces a regulator of Notch signaling (Figure 6). In GTEx Consortium data, there was correspondence between the SNP alleles associated with delayed onset and *cis*-eQTL alleles for increased expression of *CCDC82* in cortex, caudate and putamen (Figure S7). This gene also scored highly in our TWAS (p = 5.56E-05) but did not quite achieve genome-wide significance. No similar relationship with regulation of expression was seen for either *MAML2* or *JRKL*, suggesting that increased *CCDC82* expression in brain acts to delay HD onset. Relatively little is known concerning the function of CCDC82 beyond a single report that it is one of the substrates for ATM-dependent ^phosphorylation in response to H_2_O_2_ treatment (Kozlov et al., 2016). Interestingly, reduced^ expression of ATM is reported to protect against the toxicity of mutant huntingtin fragments (Lu et al., 2014).

**Figure 6.**
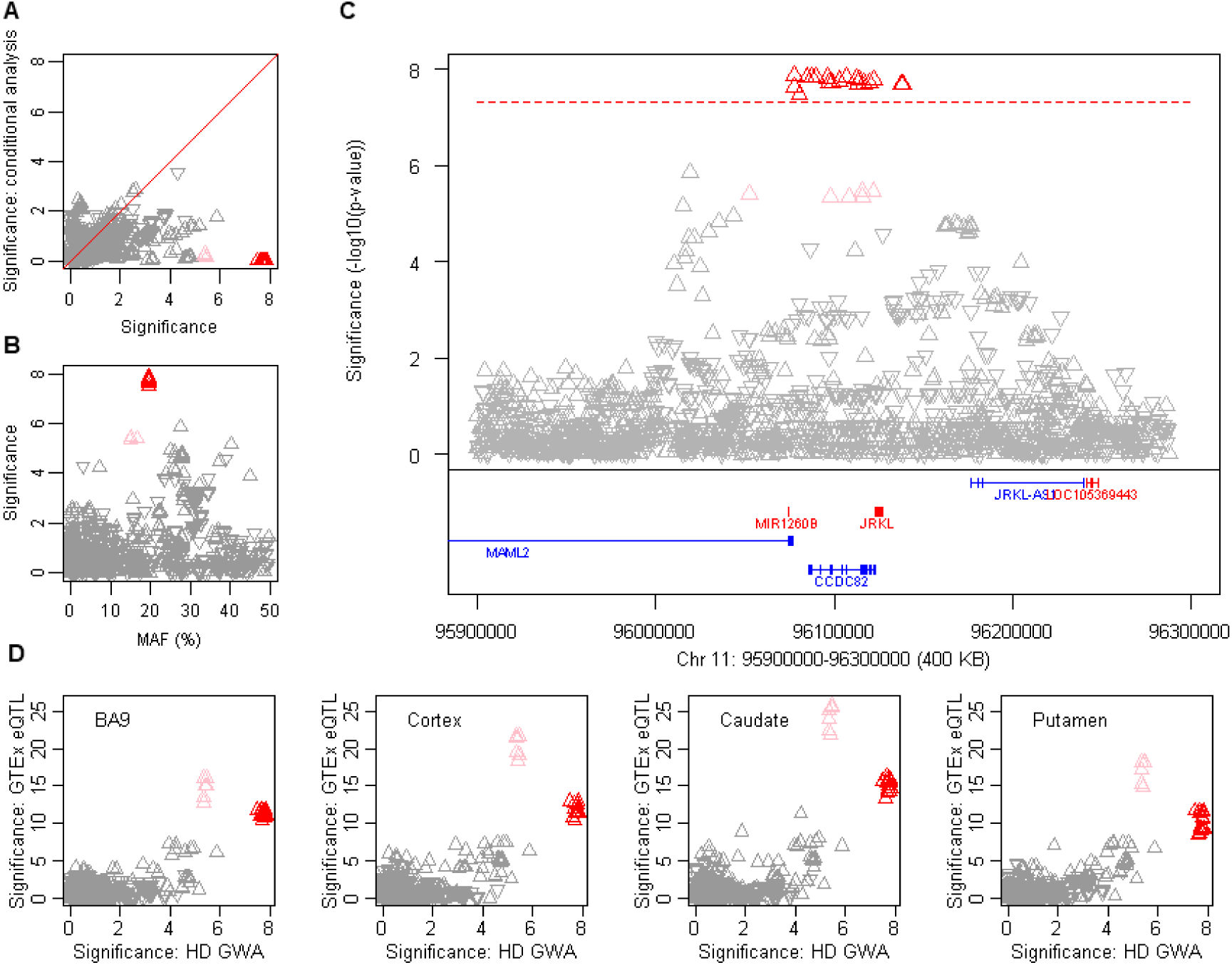
An onset modification signal on chromosome 11. **A**. Significance of SNP association in dichotomous analysis conditioned on the top SNP in the region (rs7936234, onset-delaying 11AM1, red upward triangles) is compared to GWA12345 significance, revealing the absence of independent modifier haplotypes with greater than suggestive significance (p-value < 1E-5) but the presence (with subsequent analyses) of a related haplotype (pink triangles). **B**. The significance of SNP association in GWA12345 is compared to minor allele frequency, revealing a slight difference for 11AM1 tag SNPs and those marking the closely related haplotype. **C**. Significance for SNPs is shown relative to genes in the region. **D.** Significance of SNP association in dichotomous analysis is compared to *cis*-eQTL signals for *CCDC82* in GTEx consortium data for prefrontal cortex BA9, cortex, caudate, and putamen, with the direction of the triangle indicating association of the minor allele with increased (upward) or decreased (downward) expression. Both related haplotypes (red and pink) show correlation with *cis*-eQTL SNPs associated with increased expression of *CCDC82* in all 4 brain regions.

Perhaps most intriguing is a second locus on chr 5, where a single SNP, rs79727797, supported by both continuous and dichotomous analysis, resides within an intron of *TCERG1* and near *GPR151*. *TCERG1* encodes a nuclear regulator of transcriptional elongation and pre-mRNA splicing and *GPR151* specifies an orphan G-protein receptor from the same subfamily as the somatostatin, opioid, galanin, and kisspeptin receptors. Loss-of-function variants in *GPR151* protect against obesity (Emdin et al., 2018). This region was previously implicated as potentially harboring an HD onset modifier in candidate studies of polymorphic variation of a (Gln-Ala)_n_- encoding repeat in *TCERG1*, prompted by the transcription elongation regulator 1 protein’s physical interaction with huntingtin (Holbert et al., 2001). The genome-wide significant signal associated with delayed onset detected by rs79727797 suggests that more detailed study of this locus is warranted.

## DISCUSSION

Our current GWAS findings concerning the effect of the uninterrupted *HTT* CAG repeat and the influence of DNA maintenance processes indicate that the primary target for disease onset modification is the mutation itself, rather than downstream pathogenic processes. They support a model of HD in which the rate at which disease manifestations emerge, leading to clinical diagnosis, is determined not by polyglutamine toxicity, but by length-dependent somatic expansion of the CAG repeat in critical target cells. Such somatic expansion of CAG repeats has been observed in HD and more broadly in other polyglutamine disorders (Lopez Castel et al., 2010), arguing that somatic repeat expansion may be central to the development of all polyglutamine diseases, and potentially to other repeat diseases where the causative repeat expansion lies outside the coding sequence of the gene (Nelson et al., 2013). For example, CAG expansion has been demonstrated in DNA from the cerebral cortex of post-mortem HD brains (Swami et al., 2009). Individuals with the earliest onset for any inherited CAG repeat length also showed the greatest somatic CAG expansions, consistent with somatic expansion driving the rate of the disease process. Similar to our observations of CAA interruptions in the *HTT* CAG repeat, 4 individuals with spinocerebellar ataxia 1 (SCA1) with CAT repeat interruptions of the *ATXN1* CAG repeat suggested a better tracking of age-at-onset with the uninterrupted CAG repeat size, but because these CAT codons resulted in histidine insertion, the effect was interpreted as due to the length of polyglutamine in mutant ataxin 1 rather than to somatic CAG repeat instability (Menon et al., 2013). More recently, analysis based upon the results of GWA123 showed evidence of an effect of DNA maintenance processes in modifying polyglutamine disease onset in a group of different spinocerebellar ataxias supporting the model of somatic CAG expansion as the critical driver of age-at-onset in multiple polyglutamine diseases (Bettencourt et al., 2016).

A theoretical mathematical framework has been proposed previously to explain the emergence of disease symptoms broadly in trinucleotide repeat diseases based upon somatic expansion of the underlying repeats (Kaplan et al., 2007). This computational approach argued against continuous toxicity of the mutant allele and relied instead on somatic expansion to a critical threshold repeat length as causing the dysfunction of vulnerable cells. In addition to being consistent with our GWAS findings, this mathematical model also explains the lack of earlier onset in individuals with two mutant and no normal *HTT* alleles (Figure S3B) and of similar cases in other polyglutamine diseases (Gusella and MacDonald, 2000; Lee et al., 2012b).

However, while somatic expansion of the uninterrupted CAG repeat length can explain the rate with which an HD mutation carrier develops diagnostic clinical signs, it does not speak to what causes the actual damage to vulnerable neurons once a threshold repeat size is reached. An *HTT* CAG repeat expanded above a critical threshold length might act via polyglutamine in huntingtin but it could equally act on mechanisms that have been proposed at the level of the *HTT* mRNA (Marti, 2016), splicing (Sathasivam et al., 2013), translation (Gao et al., 2017) or even via an effect at the DNA level on chromatin domains (Bruneau and Nora, 2018). The same considerations apply to the mechanisms in other CAG repeat disorders, which likely involve different threshold somatic CAG expansion lengths in different target cell types to produce disease onset. Ultimately the disease presentation in any of the trinucleotide repeat disorders may depend upon a combination of: 1) the rate at which CAG repeats expand in the critical target cell type(s), 2) the degree to which modifiers act to influence the expansion rate in that tissue, 3) the threshold somatic expansion length that must be reached for cellular damage to occur, 4) the mechanism by which that somatically expanded repeat causes the cellular damage, and 5) the degree to which the damage mechanism is influenced by modifiers. Indeed, even within a single disease these factors may come into play, since 15AM1 and 15AM2 onset-hastening and onset-delaying modifiers showed differential influences on putamen volume, total motor score and symbol digit modalities test score in premanifest HD individuals, suggesting differential effects of the modifier alleles in different target cell types critical to each phenotype (Long et al., 2018). In addition, the duration of manifest HD from onset to death is independent of CAG repeat length, suggesting either that once cellular damage has reached a critical level, HD progression is more dependent on other physiological and/or clinical factors, or that different cell types are involved in the progression of HD after onset (Keum et al., 2016). The different cell types could display different rates of CAG expansion, and potentially different threshold lengths for initiation of cellular damage, different susceptibility to modifiers, and/or different mechanisms of causing cellular damage. Indeed, the sex differences observed for some modifier effects, despite the lack of an overall sex effect on age-at-onset, argues for differential regulation in males and females of such modifier processes even in the same neuronal cell types.

The non-canonical 4AM1 and 4AM2 *HTT* haplotypes with loss or gain of CAA interruption that implicate uninterrupted CAG repeat size, and to a much lesser extent its sequence context as determinants of HD onset age, represent a very small proportion of HD chromosomes, but present both a conundrum for HD predictive testing and an opportunity for HD research. CAG repeat determination has been based largely on PCR fragment-sizing assays, but in a small percentage of individuals, a DNA sequencing assay will yield a different result with respect to uninterrupted CAG length. This difference could be particularly important for judging repeats near the borders of high normal (27-35 CAGs), reduced penetrance (36-39 CAGs) and full penetrance (> 39 CAGs) ranges and will affect molecular diagnostics. Our data also indicate that additional DNA structures will be identified on HD chromosomes as testing by DNA sequencing becomes more prevalent, further complicating interpretation. It will take yet larger sample sizes and phenotypic detail to enable prediction of outcomes from the <5% of individuals with non-canonical sequence variations, so in these cases interpretation of predictive testing should proceed cautiously. From the research perspective, additional rare *HTT* alleles with different changes in CAG sequence that currently escape ascertainment may be particularly informative. An allele with a CAA (or other codon) interruption located more centrally within a long *HTT* CAG repeat might exist without causing HD if neither CAG repeat flanking the CAA codon exceeded the 35 CAG lower threshold of the HD associated size range. This is similar to the CAT interruption scenario in SCA1 (Menon et al., 2013). Testing of this hypothesis in human HD and/or in experimental systems could support a therapeutic strategy aimed at introducing such interruptions into *HTT*.

The notion that the rate-determining driver for disease onset in HD and other repeat disorders can be modified by DNA maintenance processes also provides a prime target for developing broadly applicable therapies to prevent or delay disease onset by intervening prior to the still uncertain cellular damage mechanism(s) in each disorder. The particular DNA maintenance processes that participate in modification of HD remain to be completely defined, but include genes associated in other circumstances with ICL repair, mismatch repair and base-excision repair, among other processes. These human genes have mainly been investigated in cancer cells, where inactivating mutations have sometimes been indicted as causes of cancer initiation or progression. Interestingly, inactivation of mismatch repair genes, which causes dinucleotide repeat instability in colon cancer, suppresses trinucleotide repeat instability in mouse models, suggesting distinct mechanisms (Boland and Goel, 2010; Usdin et al., 2015). While the association with cancer for some of the HD modifier genes dictates the need for caution in manipulating these pathways to prevent neurodegenerative diseases, human genetic evidence demonstrates that the activity of the various individual genes can vary over a relatively wide range without major deleterious effects. Each of the modifier genes found here must influence HD by virtue of naturally occurring variation in either activity, expression level or regulation, yet none has emerged as a risk factor in a cancer predisposition GWAS (GWAS Catalog: https://www.ebi.ac.uk/gwas/). For modifiers where reduced activity is associated with delayed onset or increased expression is associated with hastened onset, a goal of treatment would be to reduce the activity or level of the target protein sufficiently to achieve an even greater effect than occurs due to the naturally occurring modifiers. For example, our findings with modifier 5AM1 suggest that reducing MSH3 expression levels or activity would inhibit somatic expansion of the *HTT* CAG repeat. Population data accumulated by the Exome Aggregation Consortium show that *FAN1*, *MSH3, PMS1*, *PMS2*, and *LIG1* (pLI <0.02) are all tolerant to loss-of-function (Karczewski et al., 2017) variants, which exist in the normal population at frequencies that indicate they are not being rapidly removed by selective pressure (Kosmicki et al., 2017). Even *MLH1* (pLI=0.74) is not considered to be highly intolerant of loss-of-function variants, suggesting that the activities of each of these proteins can vary over at least a 2-fold range (wild-type vs. loss-of-function heterozygote) in normal individuals without producing strong negative selection. By contrast, *HTT* (pLI =1.00) is among the most loss-of-function intolerant genes in the human genome, showing far fewer naturally occurring inactivating mutations than would be expected, presumably due to their elimination by an as yet undetermined selective pressure.

In addition to strong support for DNA maintenance processes in influencing age-at-onset of HD, the GWA12345 study also points to other potential modifier genes (e.g., *RRM2B, CCDC82*) that might modify HD by influencing the mechanism by which a somatically expanded repeat that exceeds its critical threshold precipitates cellular damage. Identification of the processes by which these genes act could provide an entrée into modifiers of disease progression. After HD clinical onset, the disease progresses over a course of ~15 years, with deterioration in motor, cognitive and, in many cases, psychiatric domains, in parallel with neurodegeneration and loss of body mass, but the duration of this manifest disease phase is independent of the inherited *HTT* CAG length. This suggests that at least some aspects of disease progression after onset involve processes or cell types different from those most important for determining age-at-onset. Indeed, the difference in significance and effect for the two top chr 5 *MSH3/DHFR* modifier haplotypes (5AM1 & 5AM2) detected using age-at-onset as the test phenotype in GWA12345 compared to the top haplotype (5AM3) detected in TRACK-HD using a multifactorial measure of progression further reinforces the potential for different modifier effects and mechanisms at different stages of disease, even shortly after onset, which is the prime period for carrying out clinical trials to delay progression of deterioration (Caron et al., 2018). The power of the GWAS approach to detect genetic modifiers that act before the onset of HD is now amply demonstrated. While the sample size can be further increased to dig deeper into this modifier pool and to support the inclusion of modifier SNPs in optimizing clinical trial power/design, the strategy should now also be applied broadly to define modifiers that influence disease progression, so a variety of approaches are being taken to define disease stages and corresponding phenotypes for GWA analysis, with the expectation that this human genetics strategy will successfully illuminate each phase of HD, from premanifest to eventual death from the disorder. This has the potential to highlight specific targets for therapeutic intervention at different disease stages, allowing a stratified approach to treatment over the protracted disease course.

## ACKNOWLEDGEMENTS

This work was supported by the CHDI Foundation, the National Institutes of Health USA (U01NS082079, R01NS091161, P50NS016367, and X01HG006074) and the Medical Research Council (UK; MR/L010305/1). T.M received a fellowship from the MRC(MR/P001629/1) and B.McA. received a studentship from the Cardiff University School of Medicine. The Enroll-HD, Registry, PHAROS, COHORT, TREND-HD, PREDICT-HD and HD-MAPS studies would not be possible without the vital contribution of the research participants and their families. Individuals who contributed to the collection subject data can be found at https://www.enroll-hd.org/acknowledgments/ for Enroll-HD, in the Supplementary Material of (Lee et al., 2017) for Registry, and in the Supplemental Information of (GeM-HD Consortium, 2015) for the GWA123 dataset.

## AUTHOR CONTRIBUTIONS

Conceptualization, J.-M.L., M.E.M., J.F.G., L.J., P.H., and S.K.; resources, R.H.M., J.P.G.V., D.L., D.K., M.O., G.B.L., J.S.P., E.R.D., I.S., and the Registry, Enroll-HD, PREDICT-HD, COHORT, PHAROS, TREND-HD, and HD-MAPS Investigators; investigation, J.L., K.A.-E., E.M.R., J.S.M., T.G., M.C., A.C., A.M.; data curation, J.-M.L., K.H.K., E.P.H., M.J.C., K.C., D.B., E.P.H., K.A.-E., T.M., B.McA., T.C.S., L.J., P.H., and M.O.; formal analysis, J.-M.L., K.C., D.B., T.G., M.J.C., P.H., C.M., T.C.S., L.H.; writing and editing: J.F.G., J.-M.L., M.E.M., V.C.W., R.M.P., K.C., T.G., J.D.L., T.M., P.H., L.J., M.C., D.G.M., S.K., C.S., A.G.E., J.S.P., E.R.D., I.S., G.B.L., and M.O.; supervision: J.-M.L., M.E.M., J.F.G., M.O., T.M., L.J., P.H., D.G.M. and S.K.

## STAR METHODS

### CONTACT FOR REAGENT AND RESOURCE SHARING

Further information and requests for resources and reagents should be directed to and will be fulfilled by the Lead Contact, James F. Gusella, Ph.D. After publication, data involving human subjects will be shared with qualified investigators given their institutional assurance that subject confidentiality will be ensured and that there will be no attempt to discover the identity of any human subject.

### EXPERIMENTAL MODEL AND SUBJECT DETAILS

Patient consents and the overall study were reviewed and approved by the Partners HealthCare Institutional Review Board. We analyzed genetic data from 9,064 HD subjects including 4,271 available from a previous GWA study (GeM-HD Consortium, 2015) and 4,793 genotyped and QC-passed in the current study. Phenotypic data for the newly analyzed subjects were made available by the ENROLL-HD platform (https://www.enroll-hd.org/) and the EHDN Registry study(http://www.ehdn.org/), who both approved our procurement of the DNA of these subjects from their repositories at BioRep Inc. (Milan, Italy). Enroll-HD is a global clinical research platform designed to facilitate clinical research in Huntington’s disease. Core datasets are collected annually from all research participants as part of this multi-center longitudinal observational study. Data are monitored for quality and accuracy using a risk-based monitoring approach. Registry is multi-centre, multi-national observational study that has been described (Orth et al., 2010). All sites are required to obtain and maintain local ethical approval.

## METHOD DETAILS

### Genome-wide SNP genotyping

DNA samples for GWA4 and GWA5 were genotyped by InfiniumOmniExpressExome-8v1-3_A (https://www.illumina.com/products/by-type/microarray-kits/infinium-omni-express-exome.html) and Multi-EthnicGlobal-8_A1 arrays (https://www.illumina.com/products/by-type/microarray-kits/infinium-multi-ethnic-global.html), respectively. Genotyping and genotype calling based on the Birdsuite algorithm (https://www.broadinstitute.org/birdsuite/birdsuite) were performed at the Broad Institute. We performed quality control (QC) analysis for each typed GWA data independently in order to generate data set for genotype imputation. Briefly, we identified HD subjects with European ancestry (based on comparison of study subjects to HapMap samples) and subsequently excluded SNPs that showed genotyping call rate < 95% or minor allele frequency < 1%.

### Genotype imputation and quality control

Each QC passed data set, including those obtained for the prior GWA1, GWA2 and GWA3, was further subjected to additional QC analyses as part of imputation process by Michigan Imputation Server (https://imputationserver.sph.umich.edu/index.html#!). Briefly, SNPs tagged due to strand mismatch and samples with low genotyping call rate in certain chunks (< 50%) were excluded. Subsequently, haplotype phasing was performed by EAGLE (v2.3.2) and genotypes were imputed using MINIMAC via the Michigan Imputation Server, using the Haplotype Reference Consortium data (Version r1.1 2016) (http://www.haplotype-reference-consortium.org) as the reference panel. Post-imputation, we further excluded SNPs with 1) imputation R square value < 0.5 in any of GWA data sets, 2) call rate < 100%, 3) Hardy-Weinberg equilibrium p-value < 1E-6 except the chromosome 4:1-5,000,000 region, or 4) minor allele frequency < 0.1%. These quality control filters generated a total of 10,986,607 imputed SNPs for 9,064 HD subjects with age-at-onset data for genetic association analysis.

### *HTT* CAG repeat genotyping assay

*HTT* uninterrupted CAG repeat size was estimated using a modified PCR amplification assay (Warner et al., 1993), adapted for the ABI3730XL DNA sequencer in a 96 well plate format. Each plate includes reactions for genomic DNA *HTT* CAG size standards, previously sequenced and known to have different uninterrupted CAG repeat lengths in the normal and expanded CAG repeat ranges. Reactions are in a total volume of 10 ul containing 1.25 mM MgCl_2_ (Applied Biosystems), 1X Buffer II (Applied Biosystems), 0.05 U Amplitaq Gold (Applied Biosystems),0.25 mM dNTPs (GE Healthcare), 1.2 ul of DMSO (Sigma), 0.125 uM each primer with 80 ng of genomic DNA. The PCR primers (forward primer labeled with 6FAM, reverse primer is tailed) are: Forward primer (HD-1) 5’ ATGAAGGCCTTCGAGTCCCTCAAGTCCTTC 3’ and Reverse primer (HD-3) 5’ GGCGGTGGCGGCTGTTGCTGCTGCTGCTGC 3’. The PCR amplification cycles are: denaturation at 94°C for 4 minutes, thirty-five cycles of denaturation at 94°C for 30 seconds, annealing at 65°C for 30 seconds, and extension at 72°C for 45 seconds, with a final extension at 72°C for 10 minutes and hold at 15°C. The PCR amplification products are loaded onto an ABI 3730XL DNA Analyzer (36 cm array, POP-7 Polymer, standard fragment analysis conditions) along with an internal size standard where 0.8 ul PCR product is loaded in 9.4 ul Hi-Di Formamide (Applied Biosystems), with 0.1 ul GeneScan 500 LIZ (Applied Biosystems). The resulting .fsa files are analyzed with GeneMapper v5.0 (Applied Biosystems) software and the CAG repeat allele sizes are estimated relative to the fragment sizes of the *HTT* CAG repeat genomic DNA standards. By convention, the CAG repeat alleles assigned to each sample are the highest peak-signal (main peak) in the normal and the expanded CAG repeat range.

### Somatic *HTT* CAG repeat expansion analysis

We classified subject blood cell DNA samples with respect to somatic CAG repeat expansion using the *HTT* CAG genotyping assay output of the GeneMapper v5.0 software (Applied Biosystems). This is not a single molecule assay but rather a bulk measure that has a floor at the inherited CAG repeat size. For a given individual, the majority of PCR products are nested around the peak signal representing the main CAG repeat size, reflecting PCR stutter inherent in the assay that masks small biological variation in CAG repeat size. However, individual samples may display PCR products at lengths detectably greater than nested bulk PCR products. These rarer products represent somatically expanded CAG repeats present in the individual. Consequently, the utility of this assay is to identify those individuals with the highest proportion of such somatic expansions, but it is not a sensitive discriminator for the majority of samples. From the GeneMapper ‘sample plot view’, a peak data table was exported for all peaks (in .txt format) containing the following information: sample name, called CAG allele, peak size in bp, peak height, area under the peak and data point/scan number of the highest point of the peak. Using the assigned main expanded CAG allele and peak size in bp data, a linear regression was performed, on a per plate basis, to assign a CAG length to all expanded peaks. Linear regression was fit using the linregress function from the SciPy Python library Stats module (http://www.scipy.org/) according to the model: *cag_i_*=β_0_ + β_1_ *size_i;_* where cag and size is the assigned main expanded CAG allele and its measured fragment size, respectively. The size data for each sample was then transformed to a CAG repeat length by using the intercept and slope from the linear regression. For peaks whose sizes transform to the same CAG length, the larger peak height was taken. The data were filtered using an upper size threshold of 500 bp and minimum peak height threshold of 50 RFU (relative fluorescent units). To calculate the proportion of expansion products for each sample, the expanded peaks were then transformed to a proportion relative to the height of the main CAG-allele assigned peak for that sample. The proportion of expansion products was then expressed as the sum of the peak-proportions, yielding the “peak proportional sum value” for that sample.

### *HTT* MiSeq DNA Sequencing

DNA sequencing of *HTT* exon 1 CAG repeat and adjacent region in genomic DNA samples was accomplished using the Illumina MiSeq Adapted Metagenomics 16S Targeted Resequencing Protocol Library Preparation guide (Part # 15044223 Rev. B) using locus specific primers. The Step 1 oligonucleotide primer pair (PCR 1) comprising the forward and reverse *HTT* CAG target sequences and Illumina Adapter sequence (in italicized text) was:

ms_hd_f

*TCGTCGGCAGCGTCAGATGTGTATAAGAGACAG*ATGAAGGCCTTCGAGTCCC

ms_hd_r

*GTCTCGTGGGCTCGGAGATGTGTATAAGAGACAG*GGCTGAGGAAGCTGAGGA

The Step 1 PCR reaction conditions in a total reaction volume of 20 ul, comprise 1X Buffer LongRange (Qiagen), 1X Q-Solution (Qiagen), 500 uM dNTP (Qiagen), 0.25 uM each forward and reverse primer, 2 units of LongRange Enzyme (Qiagen), and 20 ng of genomic DNA. PCR amplification cycles were: initial denaturation at 93°C for 3 minutes, thirty-three cycles of denaturation at 93°C for 30 seconds, annealing at 60°C for 30 seconds, and extension at 68°C for 90 seconds, and hold at 15°C until bead clean up. The Step 2 PCR reaction conditions, in a total reaction volume of 50 ul, comprise 1X Buffer LongRange (Qiagen), 1X Q-Solution (Qiagen), 500 uM dNTP (Qiagen), 0.8 uM each forward and reverse primer, 2 units of LongRange Enzyme (Qiagen), and 5 ul of Step 1 PCR product, with the following cycle parameters: initial denaturation at 93°C for 3 minutes, eight cycles of denaturation at 93°C for 30 seconds, annealing at 60°C for 30 seconds, and extension at 68°C for 90 seconds, hold at 15°C until bead clean up. The Step 2 PCR amplification products were cleaned up using a 1X bead to PCR product ratio. The libraries were pooled in sets of 24, 96, or 192 and run on the MiSeq at concentrations 8 - 15 pM with 10% PhiX spike-in of the same concentration for a 2×300 bp read. The MiSeq output in FASTQ format was utilized for analysis to determine the size of uninterrupted CAG repeat and adjacent DNA sequence.

### *HTT* uninterrupted CAG repeat length and adjacent triplet repeat DNA sequence

The length of the uninterrupted CAG repeat and the adjacent sequence were determined from #1) MiSeq, as well as from #2) whole genome sequence data. For #1) the 300 bp paired-end read MiSeq data (FASTQ format) were analyzed as follows. Each sequence read pair was processed using Python (v3.5.1). For the forward strand profiling began at the first instance of a CAGCAGCAG 9mer (5’end of the uninterrupted CAG repeat tract) and continued across each successive trinucleotide repeat until a CAGCTTCCT 9mer (3’ end of the polymorphic triplet repeat tract) was encountered or until the end of the read was reached. The read that is antisense to the forward strand was reverse-complemented and was profiled in the same manner. If the uninterrupted CAG repeat tract and adjacent triplet repeat structure matched on both mates of a read pair, then they were aggregated and counted. Unmatched forward- and reverse-strand reads were discarded. The uninterrupted CAG repeat allele was assigned (Python v3.5.1), using the distribution of uninterrupted CAG repeat sizes of the profiled structures, to construct a CAG repeat genotype for each sample by identifying the two most frequent CAG peaks. The highest frequency profiled structures that contained each uninterrupted CAG repeat allele were then used to create a complete genotype (both alleles) for the sample that encompassed both the pure CAG tract and the adjacent triplet repeat sequence. For #2), whole-genome sequence data (https://www.sfari.org/resource/simons-simplex-collection/), the CRAM files were converted to BAM file format using samtools 1.7. (Li et al., 2009). The TREDPARSEv0.7.8 software package (Tang et al., 2017) was utilized in fullsearch mode in Python v2.7.x, to assign the length of the uninterrupted CAG tract on both alleles. To determine the adjacent sequence, the reads mapping to *HTT* exon 1 were subset from the alignment (samtools 1.7). Each individual read was searched for a 9mer of CAGCTTCCT, relative to the forward strand. The DNA sequence of the read was then read in reverse (3’ to 5’) until 3 consecutive CAG repeats (CAGCAGCAG) were found (Python 3.5.1). All discovered sequences were then aggregated and counted and only those with 3 consecutive CAG repeats at the 5’ end and the 9mer at the 3’ end were taken as complete. Complete sequences that represent 14% or less of the aggregated reads were filtered out of further analysis. Samples with a single complete sequence were assigned as homozygotes and samples with two different complete sequences were assigned as heterozygotes. Based upon direct examination of the sequence reads, the length of the uninterrupted CAG repeat was that assigned by TREDPARSE for canonical chromosomes and differed by 2 for the non-canonical chromosomes based upon the use by TREDPARSE of the canonical allele in the human genome reference sequence.

## QUANTIFICATION AND STATISTICAL ANALYSIS

### Genome-wide association study (GWAS) analysis using the continuous phenotype in meta-analysis and combined analysis

For each study subject, age-at-onset of diagnostic motor signs and CAG repeat size based on the genotyping assay were used to calculate residual age-at-onset, representing years of deviation from the expectation. For example, a HD subject with a residual age-at-onset of +5 indicates an individual who developed motor symptoms 5 years later than expected (compared to the majority of HD subjects) considering their CAG repeat length. We primarily analyzed HD subjects carrying 40-55 CAG repeats to minimize the levels of inaccuracy in calculating the residual age-at-onset. In addition, dichotomized phenotype analysis was also performed to detect statistical artifacts (see next section). To determine whether our previous association analysis using GWA123 data were replicated, we performed combined analysis of the GWA4 and GWA5 data sets. The residual age-at-onset of motor symptoms data as the continuous dependent variable was modeled as a function of minor allele count of the test SNP, sex, and the first 4 principal component values from the genetic ancestry analysis in a linear mixed effect model with relationship matrix using GEMMA (version, 0.94 beta) (http://www.xzlab.org/software.html). Subsequently, we performed meta-analysis to summarize individual GWA analysis results. Each HD GWA study (i.e., GWA1, 2, 3, 4, and 5) was independently analyzed in a linear mixed effect model to test the association of SNPs with sex and genetic ancestry covariates. Then, the 5 resultant sets were combined by meta-analysis using METAL (2011-03-25 release) (https://genome.sph.umich.edu/wiki/METAL_Documentation). Finally, for the combined GWA12345 dataset, residual age-at-onset as the continuous dependent variable was modeled as a function of minor allele of the test SNP, sex, source of GWA, and first 4 principal component values from the genetic ancestry analysis in a linear mixed effect models with relationship matrix. Results of this combined continuous phenotype analysis served as the basis for the estimation of effect sizes and significances of SNPs.

### Genome-wide association study (GWAS) analysis using the dichotomous phenotype

To confirm the lack of statistical artifacts in our standard combined continuous analysis and to reveal significant SNPs that are associated with our phenotype in a non-continuous manner, we additionally performed GWA analysis using a dichotomized phenotype. Subjects were sorted based on residual age-at-onset, those with the top 30% (2,719 subjects) and bottom 30% (2,719 subjects) were chosen and assigned to phenotype groups (‘late’ and ‘early’ onset, respectively). Based on the previous analysis of GWA123, a 30% cut-off provided the best ability to detect both the common and rare modifier effects at the chr 15 locus and so was used for this GWA12345 analysis. The dichotomous phenotype data were modeled as a function of minor allele count of test SNP, sex, source of GWA study, and first 4 principal components from the genetic ancestry analysis in a fixed effect model.

### Conditional analysis

For selected candidate regions with significant association signals in the GWA12345 combined analysis using either continuous or dichotomous phenotype, we performed conditional analysis to characterize the number of independent modifier haplotypes. Having established a robust modifier effect at a locus, we then sought additional modifier haplotypes among the SNPs with suggestive signal in the overall analysis (p <1E-5). Briefly, the same statistical model for single SNP analysis with an additional covariate of the minor allele count of the top SNP in the region was constructed in a fixed effect linear model to test independence of SNPs in the region. If significant association signals remained in the first conditional analysis, the top SNP in the conditional analysis was used for the next conditional analysis to confirm independence. Subsequently, SNPs tagging each independent modifier haplotype were identified based on continuous phenotype analysis results, dichotomous phenotype analysis results, conditional analysis, and eQTL data.

### Modifier haplotypes and tagging SNPs on chromosome 4

To characterize modifier haplotypes on chromosome 4, we compared continuous phenotype analysis and conditional analysis results. When the single SNP association analysis model included the top SNP in the region (i.e., rs764154313), a group of SNPs became non-significant while others remained genome-wide significant, suggesting that two modifier haplotypes exist in this region. We defined the modifier haplotype 4AM1 tagging SNPs as SNPs that were genome-wide significant in the combined continuous phenotype analysis, not genome-wide significant in the conditional analysis using rs764154313 and associated with onset delaying effects for the minor allele (red triangles in Figure 2). SNPs that were genome-wide significant (p-value < 5E-8) in the combined continuous phenotype analysis, not genome-wide significant in the conditional analysis using rs183415333 and associated with onset hastening effects for the minor allele were assigned as modifier haplotype 4AM2 tagging SNPs (green triangles in Figure 2).

### Association analysis using residual age-at-onset based on uninterrupted CAG repeat size

For samples with atypical *HTT* CAG repeat sequences (e.g., loss of CAA interruption, duplicated CAA interruption), we re-calculated residual age-at-onset. The length of each HD subject with an atypical CAG repeat sequence was used as independent variable for our CAG-onset phenotyping model (Lee et al., 2012b). Then, re-calculated residual age-at-onset of motor symptoms was modeled as a function of the minor allele count of SNP, sex, source of GWA and genetic ancestry covariates in a fixed effect model. Overall association signals using residual age-at-onset based on genotyping CAG and those based on uninterrupted CAG repeat size were highly similar, except for the chromosome 4 region.

### Modifier haplotypes and tagging SNPs at *MSH3/DHFR*

Comparison of continuous phenotype analysis and conditional analysis results revealed 3 modifier haplotypes at the locus at *MSH3/DHFR* on chromosome 5. The first haplotype, 5AM1, was defined by the top genome-wide significant SNP, rs701383, and further 5AM1 tag SNPs were those associated with hastened onset at p<1E-5 in the continuous analysis and p>1E-5 in analysis conditioned on rs701383 (red downward triangles in Figure 3). Haplotype 5AM2 was defined by rs113361582, which remained genome-wide significant in the analysis conditioned on rs701383. Further 5AM2 tag SNPs were those associated with delayed onset at p<1E-3 in both the continuous analysis and analysis conditioned on rs701383 with minor allele frequency < 5% (green upward triangles in Figure 3). Haplotype 5AM3 was defined by rs1650742 which, at higher allele frequency than 5AM2 tag SNPs, was associated with an onset delaying signal (p<2E-6) that was not reduced by conditioning on rs113361582. Further 5AM3 tag SNPs were those with minor allele frequency between 20 and 35% associated with delayed onset at p<1E-3 in the continuous analysis, with p >1E-3 in the conditional analysis using rs701383 (purple upward triangles in Figure 3).

### Modifier haplotypes and tagging SNPs on chromosome 15

From association signals on chromosome 15, we identified 4 modifier haplotypes. Haplotype 15AM1 was marked by the top overall SNP, rs150393409, and additional 15AM1 tag SNPs were those with p<5E-8 in the combined continuous phenotype analysis, p>5E-8 in the conditional analysis using rs150393409, and p<5E-8 in the conditional analysis using rs35811129, with minor allele frequency <5% associated with hastened onset (red downward triangles in Figure 4). Haplotype 15AM2 was marked by rs35811129 and additional 15AM2 tag SNPs were those with p<5E-8 in the combined continuous analysis, p<5E-8 in the conditional analysis using rs150393409, p>5E-8 in the conditional analysis using rs35811129, with minor allele frequency >20% associated with delayed onset (green upward triangles in Figure 4). Haplotype 15AM3 was marked by rs151322829 which remained genome-wide significant in the above conditional analyses. Additional 15AM3 tag SNPs were those with p<1E-5 in the combined continuous analysis and in conditional analyses with either rs150393409 or rs35811129, with minor allele frequency <3% associated with hastened onset (purple downward triangles in Figure 4). Finally, haplotype 15AM4 was revealed by rs34017474, whose significance increased in the conditional analysis using rs150393409, and additional 15AM4 tag SNPs were those with p<1E-5 in the continuous analysis, p<5E-8 in the conditional analysis using rs150393409, and minor allele frequency >30% (gold triangles in Figure 4).

### Modifier haplotypes and tagging SNPs on chromosome 19

We identified 3 modifier haplotypes at the chromosome 19 locus. Haplotype 19AM1 was marked by the top genome-wide significant SNP, rs274883, and additional 19AM1 tag SNPs were those with p<1E-4 in the combined continuous analysis, p>1E-2 in the conditional analysis using rs274883 and associated with delayed onset (red upward triangles in Figure 5). Haplotype 19AM2 was marked by rs3730945 whose suggestive signal was not reduced by conditional analysis using rs274883. Additional 19AM2 tag SNPs were those with p<1E-5 in the combined continuous analysis, p>1E-5 in the conditional analysis using rs3730945, and minor allele frequency >30%, associated with hastened onset (green downward triangles in Figure 5). Haplotype 19AM3 was revealed by rs145821638, whose suggestive signal was not reduced by conditional analysis using rs274883, and additional 19AM3 tag SNPs were those with p<1E-2 in the combined continuous analysis, p>1E-2 in the conditional analysis using rs145821638, and minor allele frequency <2% associated with delayed onset (purple upward triangles in Figure 5).

### Transcriptome-wide association study analysis

The association of gene expression and residual age-at-onset TWAS was performed using the FUSION package (Gusev et al., 2016), imputing the Common Mind Consortium prefrontal cortex expression to the GWA12345 summary association statistics, for the 5419 genes for which there was a significant genetic component of expression. For genes with a significant TWAS association (genome-wide significance p<9.23E-6), the association between SNPs in the region and residual age-at-onset was calculated conditional on expression using FUSION. The proportion of the genetic liability to age-at-onset attributable to expression was quantified by using HESS to calculate regional heritability using the GWAS summary statistics before and after conditioning on expression (Shi et al., 2016). The ratio of the regional heritability after conditioning on expression to that before conditioning was regarded as an estimate of the proportion of genetic liability that is not attributable to expression (thus, 1 minus the ratio gives the proportion of liability that is attributable to expression).

### Pathway analysis

Gene-wide association analyses were carried out in MAGMA (de Leeuw et al., 2015) on summary statistics from GWA12345, using genotypes from GWA345 as a reference panel to estimate LD. For primary analysis, a window of 35kb upstream and 10kb downstream of gene positions (GRCh37/hg19) was used with the “multi” analysis option, combining the mean SNP p-value with the top SNP p-value (corrected for number of SNPs, LD). Pathway enrichment analyses were performed in MAGMA, correcting for LD between genes, SNPs, initially using “self contained analysis” measuring overall association among genes in a pathway, and then using a more conservative “competitive” analysis, to compare association in genes within a pathway to those outside the pathway. ALIGATOR (Holmans et al., 2009) was used to test whether pathways contain a larger number of “significant” genes (here defined as the minimum SNP p-value in that gene, chosen such that 5% of genes in the genome are considered significant), than expected by chance, given the number of SNPs they contain. Initial analysis considered the 14 pathways found to be significantly enriched for GWAS signal in GWA123 (GeM-HD Consortium, 2015) augmented by 77 DNA repair pathways taken from Pearl et al. (Pearl et al., 2015). To test for potential novel areas of disease-relevant biology, an exploratory analysis was performed on 14,210 pathways containing between 10 and 500 genes from the Gene Ontology (GO) (Gene Ontology Consortium, 2015), Kyoto Encyclopedia of Genes and Genomes (KEGG) (Kanehisa et al., 2016), Mouse Genome Informatics (MGI) (Eppig et al., 2015), Pathway Interaction Database (PID) (Schaefer et al., 2009), Protein ANalysis THrough Evolutionary Relationships (PANTHER) (Mi et al., 2013), BioCarta (Nishimura, 2001) and Reactome (Fabregat et al., 2016).

### Analysis of GTEx eQTL data

For selected candidate genes, we compared modification association signals obtained from either continuous phenotype analysis or dichotomous phenotype analysis to eQTL signals of human tissues obtained from GTEx consortium. Publicly available single tissue eQTL data (version 7) (https://gtexportal.org/home/) were downloaded. GTEx eQTL analysis calculated the effect size for a SNP relative to the alternative allele. In order to make comparable data sets, the sign of effect size of an eQTL SNP whose alternative allele is major allele in the HD data set was flipped. Then, significance values (-log10(p-value)) in HD modification GWA analysis were compared to those in GTEx eQTL analysis. SNPs from these comparisons were indicated based on the directions of eQTL; a SNP whose minor allele is associated with increased or decreased expression levels of test gene is indicated with an upward or downward triangle respectively.

### Hardy Weinberg Equilibrium Test

The 7,013 GWA samples with peak proportional sum values in the CAG somatic expansion assay were rank sorted in descending order by peak proportional sum value and samples were divided into quartiles using Python v3.5.1., yielding 1753 samples in the quartile with the highest proportion of expanded PCR products. Chi square test for deviation of SNP rs701383 minor allele count from Hardy Weinberg Equilibrium in this top quartile was performed according to the standard method using the formula: (Obs-Exp)^2^/Exp, where Obs=observed, Exp=expected. The expected minor allele count for each class was calculated from the minor allele frequency of the SNP in the entire 7,013 sample data set. The minor allele frequency was 25.7 % matching the minor allele frequency reported for Europeans.

## DATA AND SOFTWARE AVAILABILITY

Original data will be made available on request after publication. Data involving human subjects will be shared with qualified investigators given their institutional assurance that subject confidentiality will be ensured and that there will be no attempt to discover the identity of any human subject.

## SUPPLEMENTAL INFORMATION

**Table S1 – SNPs showing suggestive significance in GWA12345 – related to Figure 1 and Table 1**

All SNPs showing significance p <1E-5 in either combined continuous phenotype analysis, male-specific analysis, female-specific analysis or dichotomous phenotype analysis.

**See Table_S1.xlsx**

**Table S2 – Gene-based association – related to Figure 1 and Table 1**

Gene-wide association analyses using window of 35kb upstream and 10kb downstream of gene positions (as defined by NCBI Build 37) and no window are shown for GWA12345, GWA45, and GWA123, with genome-wide significant values, p<2.5E-6 (Kiezun et al., 2012) shown in red.

**See Table_S2_Gene-Based_Association.xlsx**

**Table S3 –Enrichment in GWA12345 of pathways from GWA123 – related to Figure 1 and Table 1**

Pathway enrichments relative to those reported in GWA123 (GeM-HD Consortium, 2015) are shown from a “self contained analysis” measuring overall association among genes in a pathway, a “competitive” analysis measuring association in genes within a pathway to those outside the pathway and analysis with ALIGATOR to test whether pathways contain a larger number of “significant” genes than expected by chance, given the number of SNPs they contain.

**See Table_S3_Enrichment_in_GWA12345_of_Pathways_from_GWA123.xlsx**

**Table S4 – Enrichment of pathways from a larger set of DNA repair pathways – related to Figure 1 and Table 1**

Pathway enrichments are shown for a larger set of DNA repair pathways (Pearl et al., 2015).

**See Table_S4_Enrichment_in_Pearl_Pathways.xlsx**

**Table S5 – Top scoring genes in pathways from Tables S3 and S4 – related to Figure 1 and Table 1**

**See Table_S5_Top_Genes_in_Enriched_Pathways_from_Tables_S3&S4.xlsx**

**Table S6 – Enrichment across 14,210 pathways**

Significant enrichments (self-contained analysis) in 14,210 pathways containing between 10 and 500 genes from the Gene Ontology (GO), Kyoto Encyclopedia of Genes and Genomes (KEGG), Mouse Genome Informatics (MGI), National Cancer Institute (NCI), Protein ANalysis THrough Evolutionary Relationships (PANTHER), BioCarta and Reactome.

**See Table_S6_Enrichment_Across_14210_pathways.xlsx**

**Table S7 – *HTT* canonical and non-canonical DNA sequence variations in HD individuals and controls** The frequencies of DNA sequence variations immediately downstream to the uninterrupted *HTT* CAG repeat on expanded (>35 CAGs) and normal (<36 CAGs) chromosomes are given, classified as canonical (CAG repeat followed by a single CAACAG codon pair) and non-canonical (CAG repeat followed by all other sequence variations). A total of 1,980 sequences were determined by MiSeq DNA sequence analysis of 990 unique HD study individuals (17 without an expanded repeat) not having either the 4AM1 or 4AM2 top SNPs. Augmenting these results, a total of 9,486 sequences were observed in a large non-HD study sample of 4,738 unique individuals, as determined by analysis of whole genome sequence data. In the total of 11,456 chromosomes (986 with expanded CAG repeats and 10,470 with normal range CAG repeats), more than 95% of both expanded and control chromosomes had the canonical single penultimate CAA interruption, followed one of twenty-six different downstream sequence variations. The great majority of these canonical chromosomes matched the reference sequence (GRCh37/hg19): CAA1 CAG1 CCG1 CCA1 CCG7 CCT2 CAGCTTCCT1, or the common polymorphic variant: CAA1 CAG1 CCG1 CCA1 CCG10 CCT2 CAGCTTCCT1. The non-canonical sequences comprised twenty-one different sequence variations. A number of canonical and non-canonical sequence variations featured codon differences that alter the proline-rich amino acid sequence that follows the polyglutamine tract. On canonical chromosomes, two infrequent sequence variations exhibited a TCG serine or a ACG threonine residue amidst the proline codons. On four rare non-canonical sequence variations the CAG repeat was followed by a CAC histidine codon. One of these also featured a CGC arginine codon. Yet another non-canonical sequence displayed a CTA leucine codon and an ACG threonine codon. Neither frame-shift codons nor stop codons were observed in this large set of chromosomes.

**See Table_S7_HTT_DNA sequence_variations.xlsx**

---

**Figure S1.**
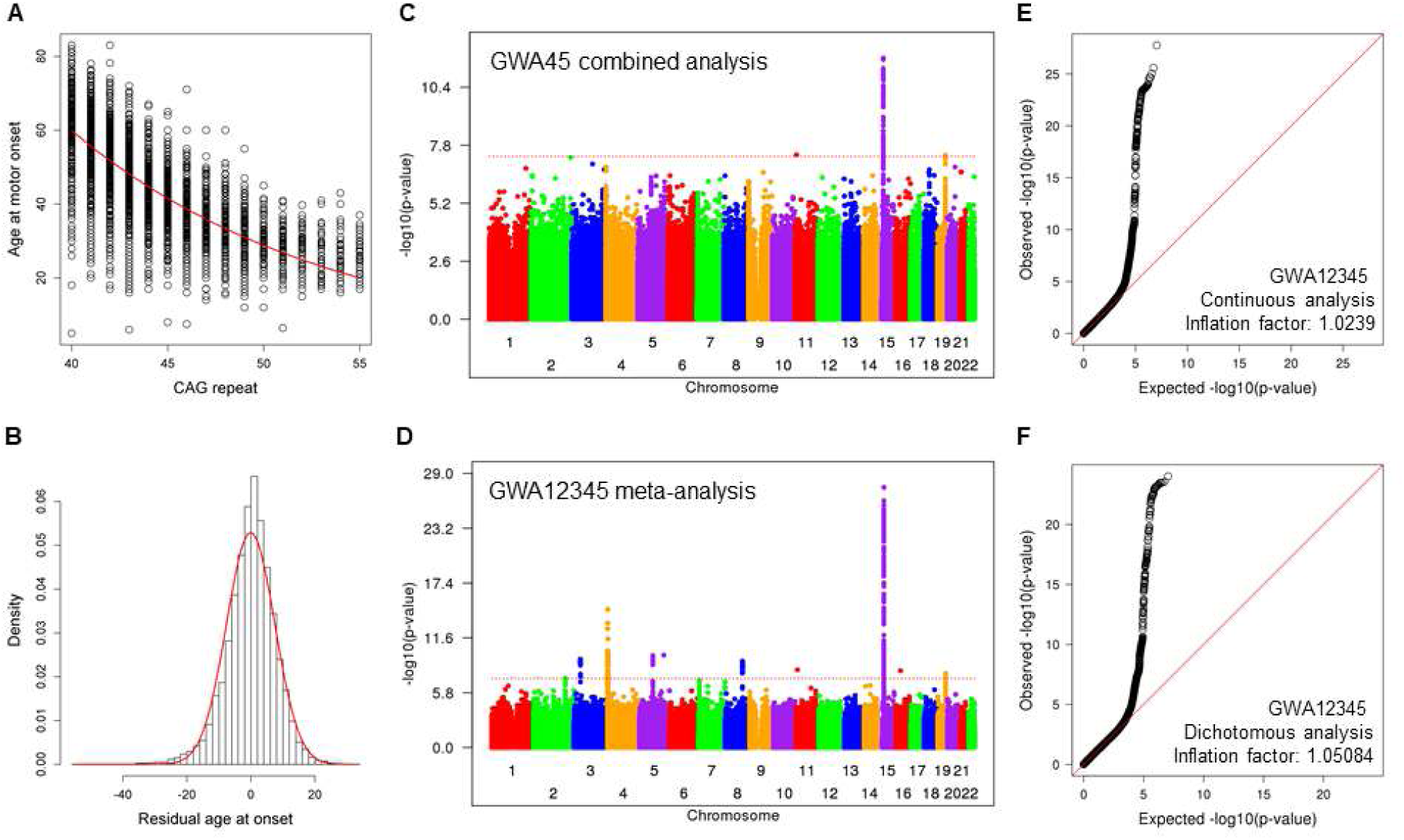
Residual age-at-onset phenotype for GWA analysis to identify genetic modifiers of HD –related to Figure 1. **A**. Age-at-onset data for individuals with HD (Y-axis) were compared to CAG repeat size based on genotyping. A red line represents our standard CAG-onset phenotype model. Residual age-at-onset was calculated for each subject by subtracting expected age-at-onset based on our CAG-onset model from observed age at motor onset. **B.** Residual age-at-onset was our primary phenotype for genetic analysis. Distribution of residual age-at-onset of individuals with HD (histogram) was compared to a theoretical normal distribution based on the mean and standard deviation of actual data to confirm data normality. **C.** For an independent comparison with our previous GWA results (GWA123), we performed mixed effect model GWA analysis of additional HD individuals (GWA45). The Manhattan plot summarizes association signals using residual age-at-onset for 4,793 HD individuals. Y- and X-axis represent-log10(p-value) and chromosome number. **D.** In addition, we performed and subsequently compared 1) meta-analysis to summarize 5 independent GWA analysis results and 2) mixed effect model combined analysis to avoid statistical artifacts. The Manhattan plot of the meta-analysis shown here and that of combined analysis using the continuous phenotype in Figure 1 were very similar, confirming the lack of batch effect in our GWA analysis results. In order to provide meaningful effect sizes of associated loci, we primarily used the results of the combined analysis. **E.** A quantile-quantile (QQ) plot based on the GWA analysis results using continuous phenotype (mixed effect model, combined analysis) and the inflation factor confirmed the lack of statistical inflation in our results. **F.** A QQ plot based on the GWA analysis results using dichotomous phenotype (fixed effect model, combined analysis) and the inflation factor confirmed the lack of statistical inflation in our results.

**Figure S2.**
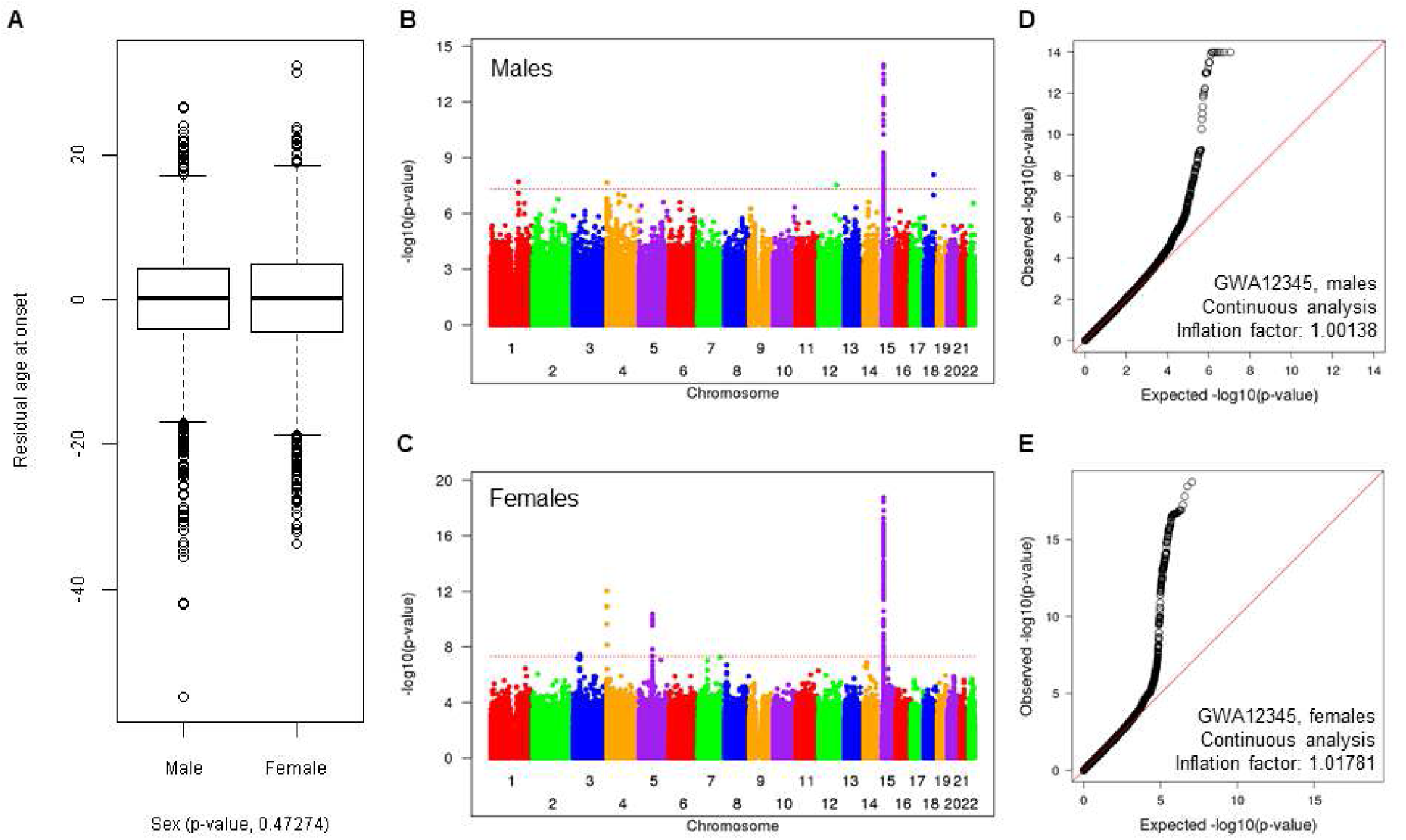
Sex-specific association analysis of residual age-at-onset – related to Figure 1. **A.** The distribution of age-at-onset residuals in GWA12345 subjects is presented as a standard box plot by sex, showing no significant difference between males and females. **B.** The Manhattan plot summarizes association signals using residual age-at-onset for 4,417 male HD individuals. Y- and X-axis represent-log10(p-value) and chromosome number. **C.** The Manhattan plot summarizes association signals using residual age-at-onset for 4,647 female HD individuals. **D.** A QQ plot based on the male-specific GWA analysis is shown, confirming lack of statistical inflation. **E.** A QQ plot based on the female-specific GWA analysis is shown, confirming lack of statistical inflation.

**Figure S3.**
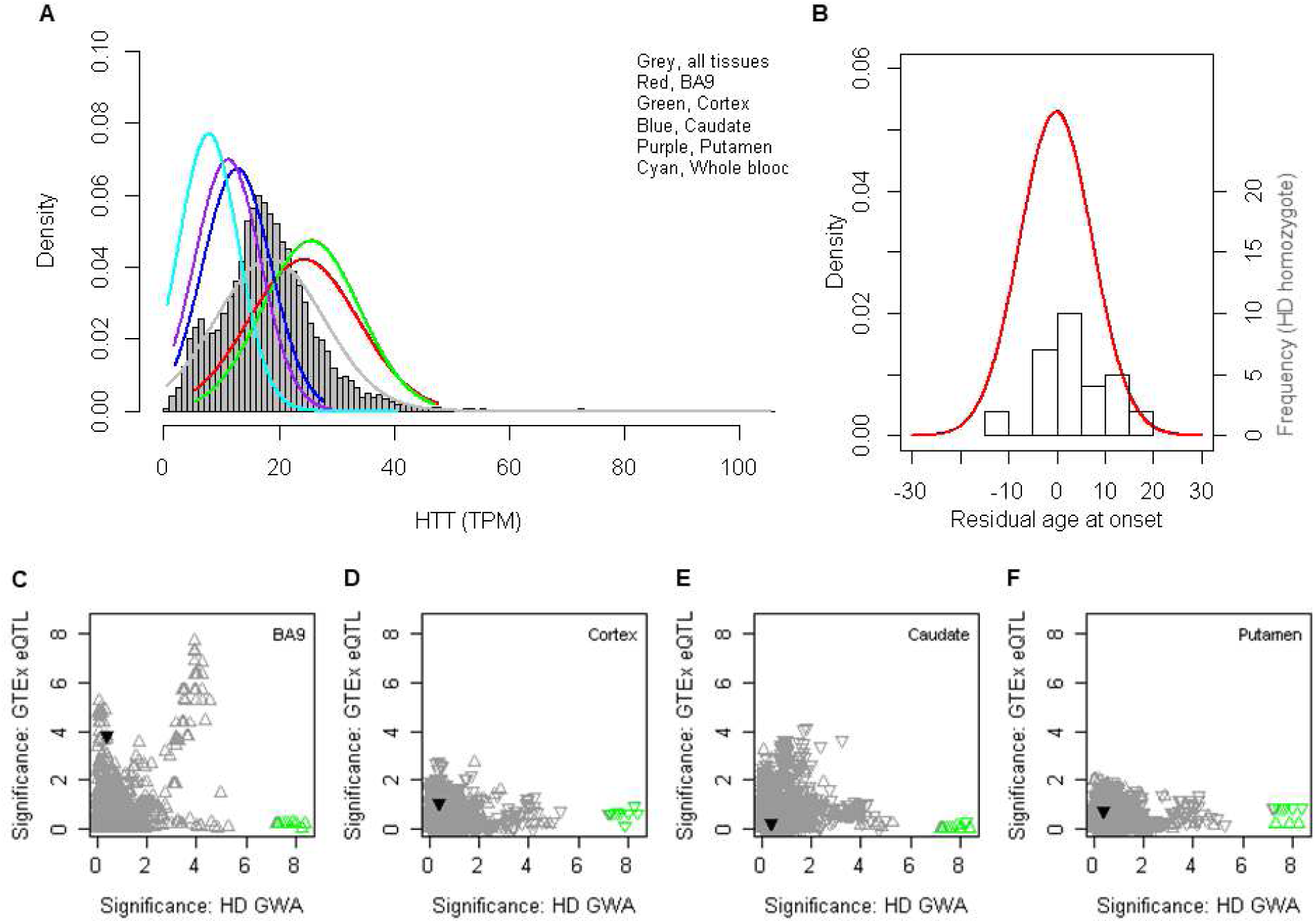
Chromosome 4 onset modifier signals vs. *cis*-eQTL signals for *HTT* – related to Figure 2. **A.** Expression levels of GTEx subjects in various tissues are plotted. The background histogram represents the distribution of *HTT* mRNA levels across all tissues. TPM (X-axis) represents Transcripts Per Million (Li et al., 2010). **B.** Residual age-at-onset of heterozygous HD individuals (red line; primary Y-axis) is qualitatively compared to a histogram distribution of residual age-at-onset of HD individuals with 2 expanded alleles (30 individuals; secondary Y-axis). **C.-F.** Onset modifier signals for chromosome 4 (X-axis) were compared to GTEx eQTL analysis results for prefrontal cortex BA9 (C), cortex (D), caudate (E), and putamen (F). Upward and downward triangles represent SNPs whose minor alleles were associated with increased and decreased *HTT* mRNA levels, respectively, in GTEx data. The 4AM1 tag SNPs are infrequent and therefore were filtered out from publicly available GTEx eQTL results. Green triangles represent tag SNPs for the modifier haplotype 4AM2 which carries a second CAA interruption but do not correspond with eQTLs in any of the brain tissues. Some 4AM2 tag SNPs are in GTEx eQTL data set, and compared to HD modifier GWA data. In BA9, GWA signals in the 1E-4 range contributed by more frequent SNPs correspond with eQTL signals for increased *HTT* expression, but these GWA signals are removed by conditioning for the 4AM2 top SNP. Notably, the 4AM2 haplotype represents a small subset (~10%) of the chromosomes that bear the more frequent rs13102260 promotor SNP (black triangle here and in Figure 2A), which was proposed to be involved in regulation of *HTT* expression levels and subsequent modification of HD (Becanovic et al., 2015) but which failed to yield a strong association signal in either GWA123 or GWA12345. Based on the ancestry of some 4AM2 HD individuals, this onset-delaying haplotype may correspond to that responsible for a Danish HD family reported previously to display later than expected onset (Norremolle et al., 2009).

**Figure S4.**
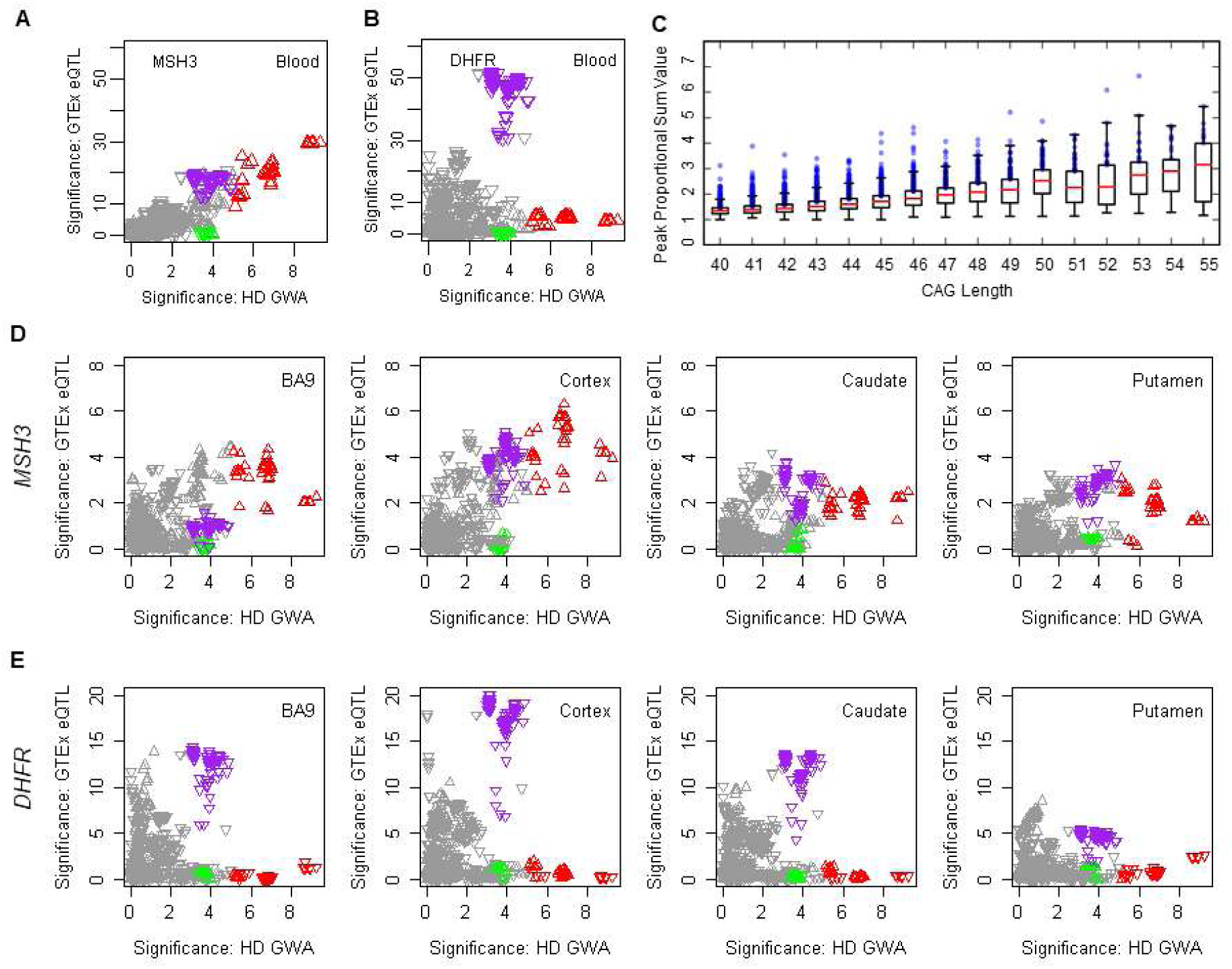
Correspondence between onset modifier signals and GTEx eQTL signals for *MSH3* and *DHFR* – related to Figure 3. In panels **A, B, D** and **E**, upward and downward triangles represent SNPs whose minor alleles were associated with increased and decreased expression levels of the test gene, respectively. Red, green, and purple triangles represent SNPs tagging modifier haplotypes 5AM1, 5AM2, and 5AM3, respectively, as in Figure 3. HD modifier GWA signals in the *MSH3*/*DHFR* region of chromosome 5 region (X-axis, -log10(p-value)) were compared to GTEx eQTL signals for *MSH3* and *DHFR* in whole blood (**A** and **B**, respectively) and in BA9, cortex, caudate, and putamen (**D** and **E**, respectively). There was no sex difference in the expression of *MSH3* in any of the tissues (blood p=0.555; BA9 p=0.586; cortex p=0.993; putamen p=0.542; or caudate p=0.054). Interaction between sex and the top 5AM1 SNP in influencing expression was not nominally significant in any tissue (blood p=0.799; BA9 p=0.867; cortex p=0.498; putamen p=0.985) except caudate (p=0.025) but the latter significance was eliminated by multiple testing correction. Larger samples sizes will be required for a definitive conclusion. **C.** The box plot shows the peak proportional sum *HTT* CAG repeat expansion values determined for 7,013 GWA12345 subjects, plotted as quartiles (whiskers1.5*IQR) for each CAG repeat size, illustrating that proportion of expanded alleles increases with CAG size and that the top quartile representing individuals with the highest proportion of somatic expansions is distinguished by a wider range of values than the other three across the CAG lengths. The values for the 1,753 individuals in the highest expansion quartile are plotted as blue circles. These individuals had 5AM1-tag SNP rs701383 deviating from Hardy-Weinberg expectation (chi-square 15.80, 2 d.f., p < 0.0004) due to an excess of the A minor allele, associated with hastened onset and increased *MSH3* expression.

**Figure S5.**
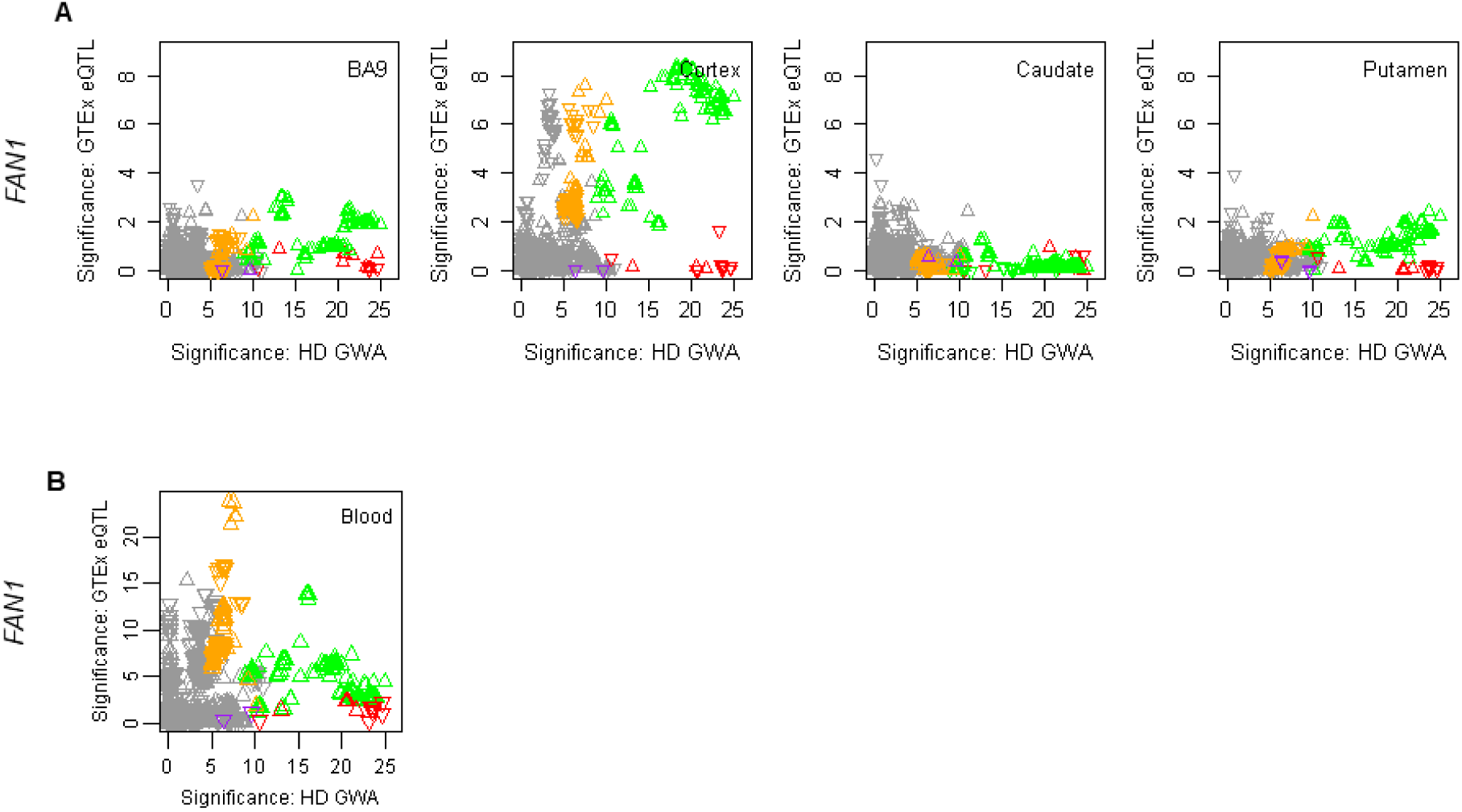
Correspondence between onset modifier signals and GTEx eQTL signals for *FAN1 –*related to Figure 4. Chromosome 15 onset modifier signals depicted (as in Figure 4) in red (15AM1), green (15AM2), purple (15AM3), and gold (15AM4) (X-axis) were compared to GTEx eQTL signals (Y-axis) for brain regions (**A**) and whole blood (**B**). Upward and downward triangles represent SNPs whose minor alleles were associated with increased and decreased expression levels of *FAN1*. Because SNPs tagging 15AM4 have alleles of close to equal frequency, the direction of the arrow varies depending on whether the minor or major allele is on the 15AM4 haplotype.

**Figure S6.**
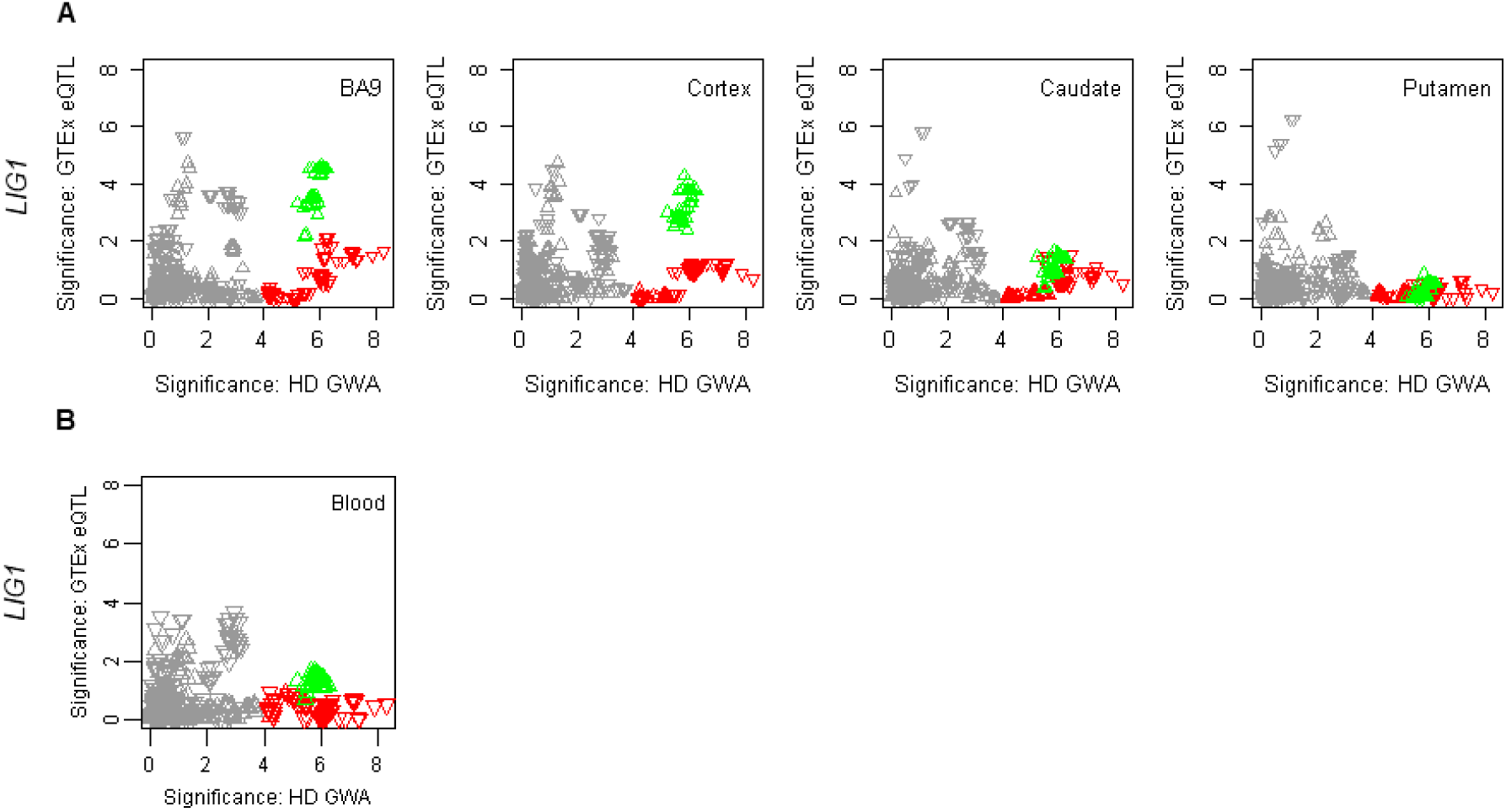
Correspondence between onset modifier signals and GTEx eQTL signals for *LIG1* – related to Figure 5. Chromosome 19 onset modifier signals depicted in red (19AM1) and green (19AM2) (X-axis) were compared to GTEx eQTL signals (Y-axis) for brain regions (**A**) and whole blood (**B**). The tag SNP for 19AM3 is too infrequent to appear in GTEx eQTL results. Upward and downward triangles represent SNPs whose minor alleles were associated with increased and decreased expression levels of *LIG1*.

**Figure S7.**
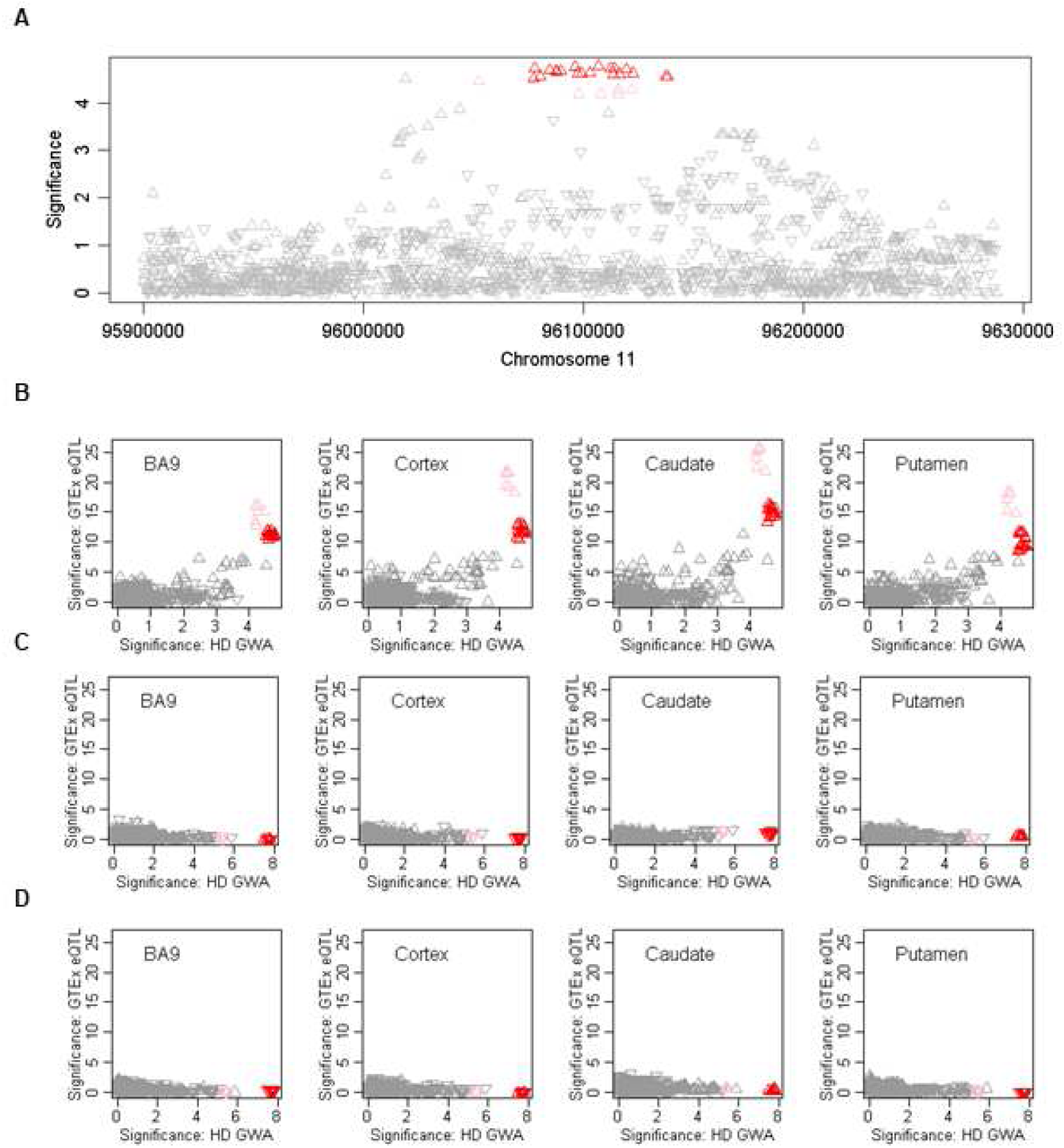
Association signals in continuous phenotype analysis for chromosome 11 region and correspondence to GTEx eQTL signals – related to Figure 6. Association analysis for SNPs in the chromosome 11 region using the continuous phenotype is shown (**A**). These continuous phenotype analysis results were compared to GTEx eQTL signals for *CCDC82* in brain regions (**B**). GTEx eQTL signals for *MAML2* (**C**) and *JRKL* (**D**) in selected brain regions (Y-axis) were compared to chromosome 11 onset modifier signals based on the dichotomous phenotype (from Figure 6). Red and pink triangles represent SNPs marking the 11AM1 and a closely related haplotype.

